# Brain-wide arousal signals are segregated from movement planning in the superior colliculus

**DOI:** 10.1101/2024.04.26.591284

**Authors:** Richard Johnston, Matthew A. Smith

## Abstract

The superior colliculus (SC) is traditionally considered a brain region that functions as an interface between processing visual inputs and generating eye movement outputs. Although its role as a primary reflex center is thought to be conserved across vertebrate species, evidence suggests that the SC has evolved to support higher-order cognitive functions including spatial attention. When it comes to oculomotor areas such as the SC, it is critical that high precision fixation and eye movements are maintained even in the presence of signals related to ongoing changes in cognition and brain state, both of which have the potential to interfere with eye position encoding and movement generation. In this study, we recorded spiking responses of neuronal populations in the SC while monkeys performed a memory-guided saccade task and found that the activity of some of the neurons fluctuated over tens of minutes. By leveraging the statistical power afforded by high-dimensional neuronal recordings, we were able to identify a low-dimensional pattern of activity that was correlated with pupil size and simultaneously recorded data in the prefrontal cortex (PFC), consistent with slow changes in the monkeys’ arousal levels while they were performing the task. Importantly, we found that the spiking responses of deep-layer SC neurons were less correlated with this brain-wide arousal signal, and that neural activity associated with changes in pupil size and saccade tuning did not overlap in population activity space with movement initiation signals. Taken together, these findings provide a framework for understanding how signals related to cognition and arousal can be embedded in the population activity of oculomotor structures without compromising the fidelity of the motor output.

## Introduction

For humans and animals to effectively interact with a fast-changing environment, they must be adept at performing a range of visually-guided motor actions. Basic everyday tasks that are critical for our survival (e.g. making an eye movement to a car that is about to cross our path) require visual information to be rapidly encoded and transformed into a motor action. In non-mammals such as birds, reptiles and fish, visually-guided motor actions (e.g. prey capture) are generated by circuits in the optic tectum (Wylie et al., 2009; Kardamakis et al., 2015; Suzuki et al., 2019; Allen et al., 2021; Isa et al., 2021) whereas in mammals such as rodents, cats and monkeys they are supported by activity in a homologous structure called the superior colliculus (SC) (Wurtz and Albano, 1980; Sparks and Mays, 1990; Gandhi and Katnani, 2011; Cang et al., 2018; Wheatcroft et al., 2022; Hafed et al., 2023).

The SC resides in the roof of the brainstem and receives input from retinal ganglion cells (Perry and Cowey, 1984; Qu et al., 2006), the primary visual cortex, and extrastriate areas such as V4 and the middle temporal area (Cerkevich et al., 2014). In addition to these visual inputs, research has shown that it also receives projections from cortical regions that have been implicated in eye movement control including the supplementary eye field (Huerta and Kaas, 1990; Parthasarathy et al., 1992) and the frontal eye field (Komatsu and Suzuki, 1985; Stanton et al., 1988; Parthasarathy et al., 1992). The role of these projections in the visuomotor transformation depends on the functional layer of the SC in which they terminate. Decades of research has shown that the SC comprises three main layers each of which contains cells with distinct neurophysiological response properties (May, 2006; Krauzlis et al., 2013; Basso and May, 2017). For example, neurons in the superficial layers, which are thought to represent the earliest stage of the visuomotor transformation, only respond to visual stimuli presented within their receptive field (RF) (Goldberg and Wurtz, 1972). In contrast, neurons in the intermediate layers elicit a response during both the presentation of visual stimuli and when saccades are made into their RF. Finally, neurons in the deep layers primarily respond in and around the time of a saccade. As is the case for neurons in the intermediate layers, they form a topographically organized map of visual space (Robinson, 1972; Katnani and Gandhi, 2011; White et al., 2017) and encode saccades across a range of directions and amplitudes (Gandhi and Katnani, 2011). Deep-layer SC neurons project to downstream nuclei innervating the extraocular muscles (e.g. the central mesencephalic reticular formation) (Sparks, 1986; Sparks and Hartwich-Young, 1989; Bohlen et al., 2017) but thresholds for evoking saccades using electrical microstimulation are often lowest in the intermediate layers (Mohler and Wurtz, 1976) suggesting that the intermediate and deep layers of the SC function together to control saccadic eye movements (Wurtz and Albano, 1980).

Although the SC is best known for its role in visually-guided motor actions, our understanding of its function has evolved considerably and recent work has demonstrated its role in processes linked to higher order cognition (Knudsen, 2011; Krauzlis et al., 2013; Clark et al., 2015; Basso et al., 2021; Rosen and Freedman, 2025). For example, behavioral performance on covert attention tasks (in which subjects are cued to attend to a stimulus presented at a specific portion of visual space) is modulated when the activity of SC neurons with RFs at the cued location are perturbed using reversible inactivation or microstimulation (Müller et al., 2005; Zénon and Krauzlis, 2012). In addition to cognitive signals, the SC is interconnected with brain regions that control arousal levels, e.g. the locus coeruleus (LC) (Edwards et al., 1979; Aston-Jones and Cohen, 2005; Sara and Bouret, 2012; Li et al., 2018; van den Brink et al., 2019; Plummer et al., 2020; Benavidez et al., 2021). Several studies using microstimulation have been able to establish a link between SC activity and non-invasive markers of arousal such as pupil size (Wang et al., 2012, 2014; Joshi et al., 2016; Wang and Munoz, 2021). Furthermore, research has shown that oculomotor behavior is impacted by changes in arousal (Di Stasi et al., 2013). For example, participants are faster at initiating visually-guided saccades to peripheral target stimuli when a startling auditory stimulus is presented prior to the onset of the target (Castellote et al., 2007). These changes, which are thought to reflect increased levels of arousal, are accompanied by increased pupil size and decreased microsaccade rate even when participants perform auditory tasks that do not necessitate saccadic eye movements (Contadini-Wright et al., 2023).

To support signals related to higher-order cognition, the SC has had to undergo a number of adaptations (Basso et al., 2021). The precise details of how conserved neuronal cell types give rise to complex processes such as spatial attention has yet to be fully elucidated. However, it must occur in a manner that does not interfere with the signals needed for volitional control of the eyes be it in the context of stable fixation or large amplitude saccades. As is the case for cortical regions that have a direct influence on motor behavior, having signals related to higher-order cognition and brain state intermingled with those used for movement encoding could be problematic as it may lead to interference with motor planning via premature or unwanted muscle contractions. In the motor cortex, research using delayed-reach tasks has shown that population responses during movement preparation are kept separate from those used during movement initiation (Kaufman et al., 2014; Elsayed et al., 2016). A similar separation has been observed for visual and motor responses in the SC (Jagadisan and Gandhi, 2022; Ayar et al., 2023; Baumann et al., 2023). For example, Jagadisan and Gandhi (2022) used linear microelectrode arrays to investigate why early eye movements are not triggered when neuronal responses to a visual target, presented before a delayed saccade to that target, cross a threshold. They found that population activity in the SC was less stable during the visual epoch of a delayed saccade task, relative to the saccade epoch. Moreover, saccades could be evoked more easily by patterned microstimulation when the temporal structure of the microstimulation was stable across electrodes, providing a potential explanation for how downstream regions differentiate between visual and motor responses. Similar results were reported by Baumann et al. (2023) who found that the strength of SC motor responses during a saccade to a visual image depends on the features of that image (e.g., contrast, orientation). When dimensionality reduction was applied to the spiking responses of neuronal populations in the SC, the population trajectory during the initial visual response to the image was orthogonal to that during the motor response. These findings replicate the separation in temporal population structure reported by Jagadisan and Gandhi (2022) and support the results of Ayar et al. (2023). They found that, although not completely orthogonal, population activity in the SC is distinct for visual and motor responses during the same oculomotor task and across different tasks, which could further facilitate the decoding of signals related to sensation, action and context by downstream regions. In light of these findings, we hypothesized that a similar mechanism might exist in the SC to prevent arousal-related fluctuations from interfering with eye movement outputs.

In this study, we observed slow fluctuations in the spiking responses of neuronal populations in the SC while monkeys performed a task that is commonly used to study visually-guided motor actions. By using dimensionality reduction on the neural population data, we were able to identify an axis in population activity space that was similar to the “slow drift axis” that has been observed in several cortical areas (Cowley et al., 2020) and is known to be associated with a constellation of arousal-related variables including pupil size and alpha power as measured using scalp electroencephalography (EEG) (Johnston et al., 2022a, 2022b). We found that this low-dimensional pattern of activity in the SC was also correlated with pupil size in the present study and with simultaneously recorded data in the prefrontal cortex (PFC), pointing to a link between this brain-wide fluctuation and changes in the monkeys’ arousal levels while performing the task. Additionally, we found evidence that this signal is isolated from SC neurons with strong saccadic responses: the activity of these cells was less correlated with the slow drift axis when it was computed using population activity in the SC and simultaneously recorded PFC data. Finally, we were able to show that signals in the SC that are used to control the size of the pupil reside in an orthogonal subspace to those that are used for saccade generation. This provides a potential mechanism through which signals related to cognition and arousal can exist in the SC, and even impact processing at the early to mid-stages of the visuomotor transformation, without leading to unwanted changes in SC neurons that are most active around the time of a saccade.

## Results

To determine how arousal-related signals related to motor-related activity in the SC, we trained two monkeys to perform a memory-guided saccade task (MGS) task (Figure 1A). During the task, subjects maintained fixation on a central point while a brief target stimulus was presented in the periphery. The subjects had to remember the location of this stimulus during a delay period and make a saccade to that location when the fixation point was extinguished. While the subjects performed the task (Monkey Do = 25 sessions, Monkey Wa = 26 sessions), we recorded the spiking responses of neuronal populations in: 1) the SC using a 16-channel linear microelectrode array; and 2) the PFC (area 8Ar) using a 96-channel “Utah” array (Figure 1B).

**Figure 1.**
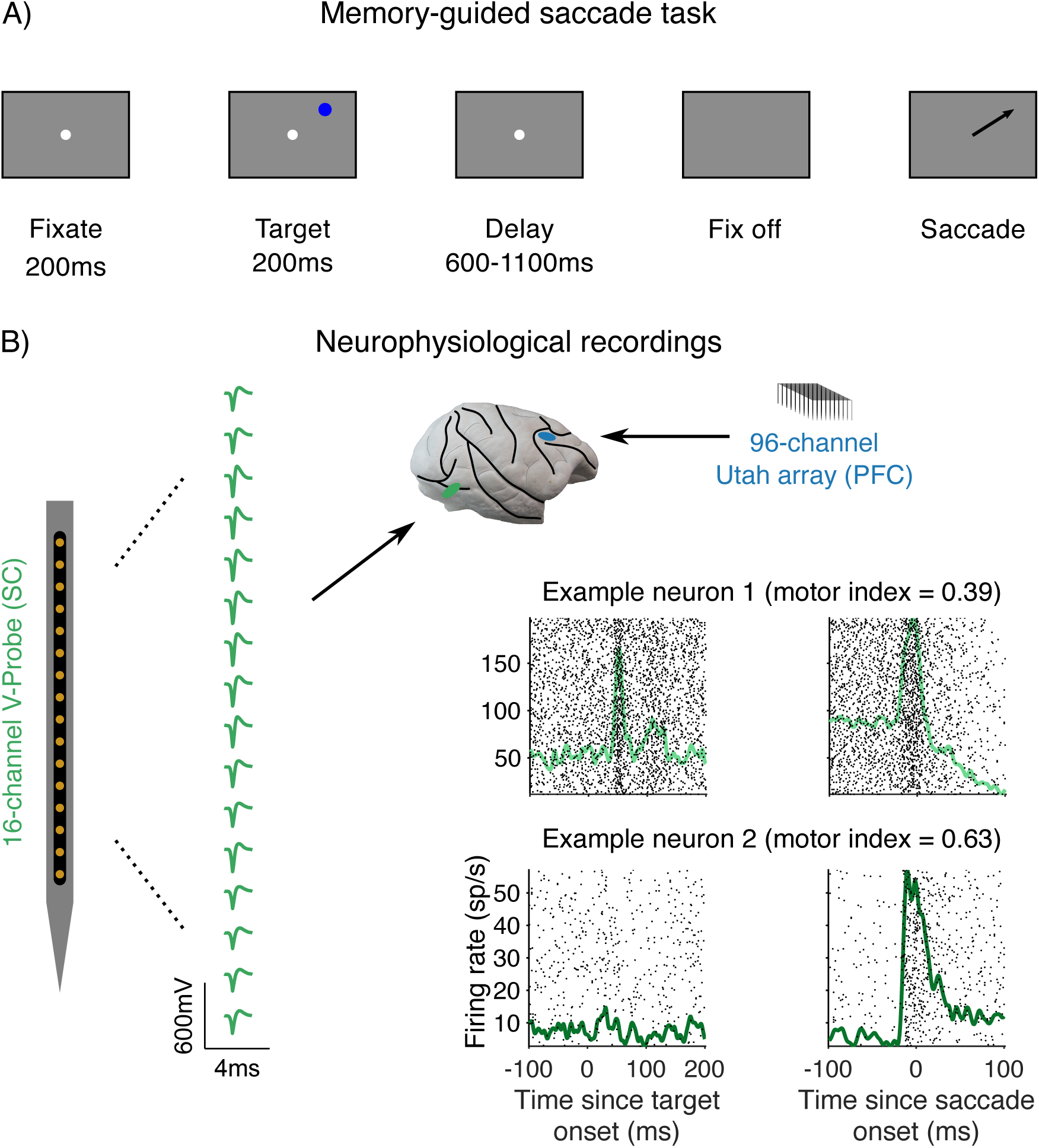
Experimental methods. (A) Memory-guided saccade task. After an initial fixation period, a target stimulus was presented at one of eight peripheral locations. This was followed by a delay period, the duration of which was chosen at random from a uniform distribution spanning 600-1100ms. The subjects’ task was to remember the location of the target stimulus and make a saccade to it when the fixation point was extinguished. (B) Neurophysiological recordings. The spiking responses of neuronal populations in the SC were recorded for targets that were visually identified to be within the neuron’s RF by applying low-amplitude microstimulation to contacts along the linear microelectrode array to evoke saccades (see Methods). The 16-channel linear microelectrode array was lowered into the brain through a recording chamber and the mean waveform for each channel during a single recording session is shown alongside a schematic of the linear array for one session. PSTHs aligned to target onset (left) and saccade onset (right) were computed to determine the response properties of individual SC neurons. Firing rates were computed in 1ms bins, averaged across trials, and smoothed using a Gaussian function for visualization purposes only (σ = 5ms).

To determine the neurophysiological response properties of individual SC neurons, we took advantage of the fact that the MGS task contains distinct epochs that can dissociate visual responses from those occurring during the execution of saccadic eye movements (Hikosaka and Wurtz, 1983) (Figure 1A). For each neuron, peri-stimulus time histograms (PSTHs) were computed relative to: 1) the onset of the target stimulus; and 2) the onset of the saccade to the remembered target location (Figure 1B). We found that some of the neurons exhibited a visual response shortly after the presentation of the stimulus and a saccadic response when an eye movement was made to the remembered target location (Figure 1B, Example neuron 1), traditionally termed visuomotor neurons, and were present mostly on the upper and middle recording contacts of our linear array. Others only exhibited a saccadic response (Figure 1B, Example neuron 2), traditionally termed motor neurons, and were present mostly on the lower recording contacts of our linear array. Based on what is known about the response properties of neurons in the SC, these findings suggest that our linear array recordings targeted neurons residing at the mid to late stages of the visuomotor transformation i.e. the intermediate and deep layers of the SC (Basso and May, 2017).

To quantify the observations we made about the strength of the saccadic response, we computed a motor index for each neuron (see Methods). SC neurons with a higher motor index (i.e. those with a stronger saccadic response relative to a baseline) were assumed to reside closer to the motor output stage of the visuomotor transformation than neurons with a lower motor index. Across sessions, we found that the mean motor index for all recorded SC neurons was 0.2761 (SD = 0.1924) (Figure 1 - figure supplement 1). Neurons recorded on deeper contacts tended to have a higher motor index in most sessions (see Figure 3A for an example), though our sampling was not equal across sessions and in some cases we found mostly visuomotor or mostly motor neurons across the responsive contacts on the linear array.

### Slow fluctuations in SC spiking activity are correlated with pupil size and PFC activity

We first asked if arousal-related fluctuations are present in the SC. As in previous studies that recorded from neurons in the cortex (Cowley et al., 2020), we found that the mean spiking responses of individual SC neurons during the delay period (chosen at random on each trial from a uniform distribution spanning 600-1100ms, see Methods) fluctuated over the course of a session while the monkeys performed the MGS task (Figure 2A, left). Note that this effect cannot be attributed to signals related to stimulus or saccade tuning as residual spike counts were computed by subtracting the mean response to each target location from trial-to-trial responses to each target location. These fluctuations are also not likely to reflect instabilities in our linear array recordings, as several steps were taken to remove neurons that exhibited changes in waveform shape over time or large sudden changes in activity level (see Methods).

**Figure 2.**
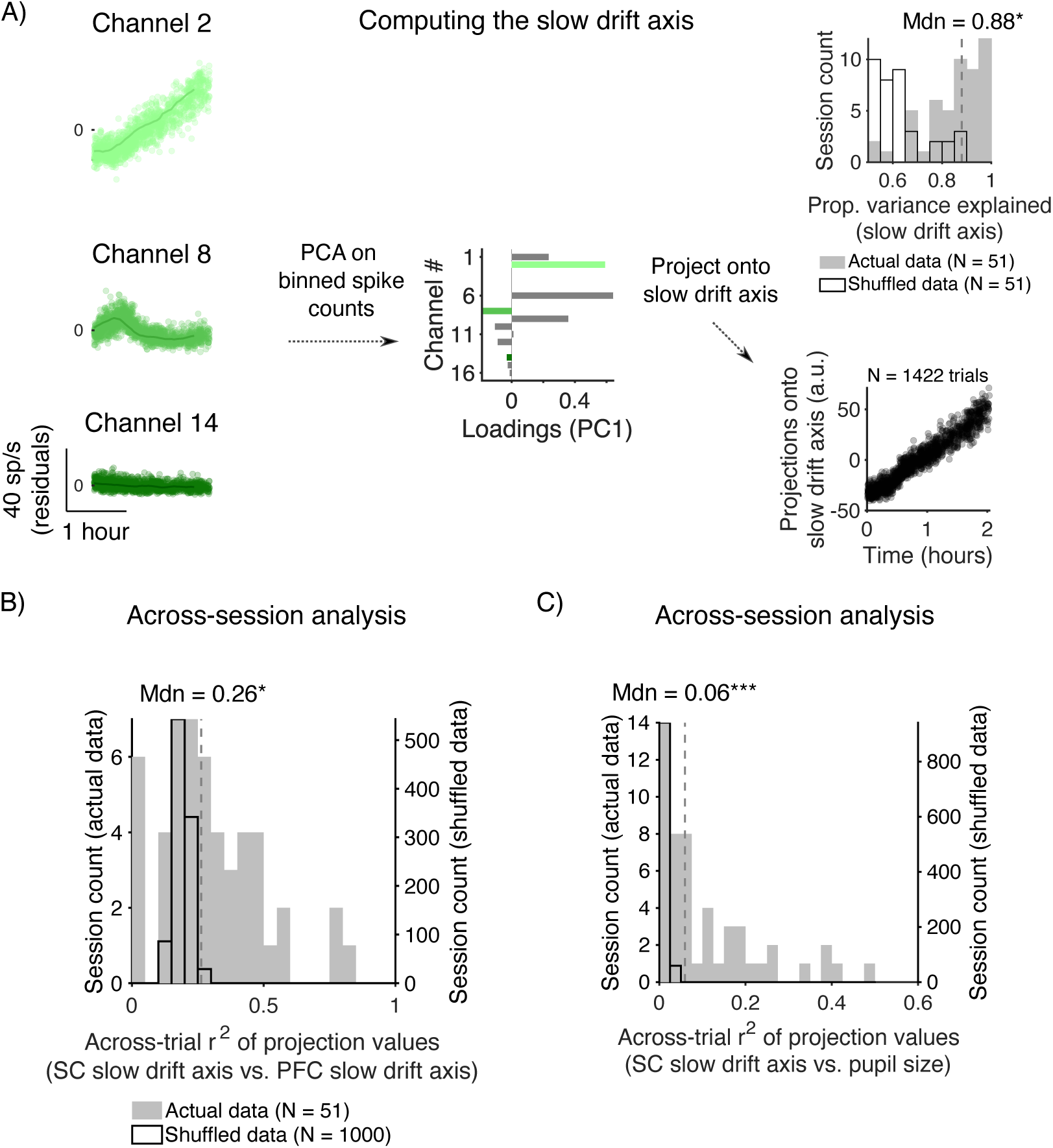
Arousal-related fluctuations are present in the SC and correlated with pupil size and fluctuations in PFC activity. (A) Computing the slow drift axis. Three example SC neurons from the same recording session (left). Each point represents the mean residual firing rate during the delay period. When PCA was applied to binned spike counts (solid line overlaid on each neuron’s spike count residuals) it yielded a vector of loadings for the first principal component, which we will term the “slow drift axis” (middle). trial-to-trial residual SC responses during the delay period were then projected onto this vector (bottom right). The histogram in the top right shows the proportion of variance explained by the slow drift axis across sessions. The median value is indicated by the vertical dashed line overlaid on the histogram. A p-value was computed by comparing the actual distribution of values to a shuffled distribution (see Methods). (B) Histogram showing the proportion of variance explained between projections onto the SC slow drift axis and projections onto the PFC slow drift axis. Note that in this analysis, the slow drift axis was computed separately for each region using the method shown in (A). Trial-to-trial residual SC responses during the delay period and trial-to-trial residual PFC responses were then projected onto the SC slow drift axis and the PFC slow drift axis, respectively. The median value across sessions is denoted by the vertical dashed line overlaid on the histogram. A p-value was computed by comparing the actual distribution of values to a shuffled distribution (see Methods). (C) Same as (B) but in this case the histogram shows the proportion of variance explained between projections onto the slow drift axis and pupil size. p < 0.05*, p < 0.01**, p < 0.001***.

To identify coordinated fluctuations in activity across the population of neurons, we applied PCA to residual spike counts of individual neurons during the delay period that had been binned using a 15-min sliding window stepped every 3 mins. This yielded a vector of loadings (Figure 2A, middle) that we termed the “slow drift axis” in line with previous studies (Cowley et al., 2020). The value of each loading corresponds to the correlation coefficient between the binned response of each SC neuron and the axis in population activity space that explained the most variance in the data. Thus, neurons whose residual activity fluctuated more over the course of a recording session had a greater portion of their variance explained by the slow drift axis (e.g. Figure 2A, Channel 2). After we identified this axis from the binned data, we then projected the residual responses of each SC neuron on single trials, which resulted in a single dimension along which we examined fluctuations over time (Figure 2A, bottom right). When the data were pooled across sessions, we found that the median proportion of variance explained by the slow drift axis was 0.88 (Figure 2A, top right). By comparing the actual distribution of values to a null distribution that was generated by shuffling the trial-to-trial spike counts for each neuron prior to performing PCA, we were able to show that the variance explained by the slow drift axis was significantly greater than what would be expected by chance (p = 0.0385). Note also that the dynamics of the SC slow drift were highly similar when it was computed using spiking responses during the baseline, visual and saccade epochs (Figure 2 - figure supplement 1). These findings suggest that arousal-related fluctuations are present throughout different task epochs and are less evident in deep-layer SC neurons independent of the time window in which spiking responses were computed.

To investigate if the SC slow drift axis was related to a global signal that might be related to shifts in arousal, we computed an identical axis using simultaneously recorded PFC data. If the SC slow drift axis represented an arousal axis that reflected coordinated activity fluctuations throughout the brain, one would expect it to be correlated with simultaneously recorded data in the PFC. For each session, we computed the correlation (Pearson’s product-moment correlation coefficient) between projections onto the SC slow drift axis and projections onto the PFC slow drift axis. As shown in Figure 2B, we found that the median variance explained across sessions, as measured using the coefficient of determination (r^2^), was 0.2627. To determine whether or not this effect was statistically significant, we compared the actual distribution of r^2^ values to a null distribution that was generated by shuffling across sessions and recomputing the correlation between projection values. We found that the proportion of variance explained was significantly greater than what would be expected if there was no correlation between projections onto the SC slow drift axis and the PFC slow drift axis (p = 0.0190). These findings suggest that slow fluctuations in responsivity are present in populations of SC neurons, and furthermore that they are coordinated across the brain. To further explore how they relate to changes in arousal or brain state, we next asked if the SC slow drift axis is associated with eye metrics that are known to be correlated with arousal levels.

Decades of research has shown that the size of the pupil can be used as a reliable marker of arousal (Mathôt, 2018; Joshi and Gold, 2020). Heightened levels of arousal are accompanied by concomitant increases in LC activity and pupil size (Aston-Jones and Cohen, 2005; Murphy et al., 2014; Joshi et al., 2016; Reimer et al., 2016; de Gee et al., 2017; Breton-Provencher and Sur, 2019). Moreover, previous work in our laboratory computed a similar slow-drift axis using spiking activity in visual cortex (V4) and PFC, and investigated the relationship between these low-dimensional neural activity patterns and different eye-related metrics (e.g., pupil size, microsaccade rate, reaction time, saccade velocity). In addition to observing a strong correlation between V4 and PFC slow drift, we found that, relative to the other eye-related metrics, pupil size was the strongest predictor of these fluctuations (Johnston et al., 2022a). Thus, to further confirm the link between the SC slow drift axis and changes in the monkeys’ arousal levels while they performed the MGS task, we next sought to explore if projections onto the SC slow drift axis were associated with pupil size. For each session, we computed the correlation (Pearson’s product-moment correlation coefficient) between projections onto the SC slow drift axis and mean pupil size during the first 200ms of the delay period when a task-related pupil response could be observed. Across sessions, we found that the median variance explained between projections onto the SC slow drift axis and pupil size was 0.0597 (Figure 2C). When the actual distribution of values was compared to an across-session shuffle distribution, we found that the proportion of variance explained was significantly greater than what would be expected if there was no correlation between SC slow drift and pupil size (p < 0.001). Interestingly, the median variance explained between projections onto the SC slow drift axis and other eye metrics including microsaccade rate was much smaller (r^2^ = 0.0026, p < 0.001) (Figure 2 - figure supplement 2). Although microsaccade rate (i.e. the number of small eye movements made during a period of steady fixation) has been linked to arousal (Siegenthaler et al., 2014; Johnston et al., 2022a; Contadini-Wright et al., 2023), microsaccade rates are task-dependent (Winterson and Collewijn, 1976; Ko et al., 2010; Bowers et al., 2021) and may be driven by activity in cortical as opposed to subcortical regions (Hafed et al., 2021). This might explain why no significant correlation was found between SC slow drift and microsaccade rate in the present study.

Taken together, these findings point to a link between the SC slow drift axis and brain-wide arousal signals. That simultaneously recorded PFC data and pupil size were correlated with projections onto the slow drift axis suggests that the signal we identified in the SC is highly similar to that which has been observed across several cortical areas (Cowley et al., 2020; Hennig et al., 2021). The fact that signals related to arousal are present in the SC is not particularly surprising given that its functional role has evolved and is now thought to include higher order cognitive processes (Knudsen, 2011; Krauzlis et al., 2013; Clark et al., 2015; Basso et al., 2021). However, it also poses a problem. Given the well-established role of deep-layers SC neurons in saccade generation (Sparks and Hartwich-Young, 1989; Gandhi and Katnani, 2011), having signals related to cognition and arousal embedded in oculomotor circuits that send gaze commands to downstream nuclei could lead to unwanted changes in eye position. One possibility is that arousal-related fluctuations are isolated from these deep-layer SC neurons that reside closer to the motor output. If this is the case, one would expect neurons with a stronger saccadic response to minimally fluctuate over time, and have a smaller portion of their variance explained by fluctuations along the slow drift axis.

### Arousal-related fluctuations are isolated from SC neurons with a higher motor index

To investigate if arousal-related fluctuations are isolated from deep-layer SC neurons, we took advantage of the fact that our linear array recordings targeted neurons residing in earlier and later stages of the visuomotor transformation (Figure 1B). As described above, this was quantified by computing a motor index for each SC neuron with high values indicating that a cell elicited a strong saccadic response to the remembered target location and low values indicating that a cell elicited a weak saccadic response. Cells with a higher motor index tend to reside in the deep layers of the SC closer to the motor output (Basso and May, 2017). In several sessions, such as that shown in Figure 3A, we observed that the loadings for the slow drift axis (i.e. the correlation between the response of each neuron and the component that explained the most variance in the data) were weaker for neurons with a higher motor index. Of note in this particular session is that these neurons were recorded from channels at the bottom of the probe, which presumably targeted the deep layers of the SC.

**Figure 3.**
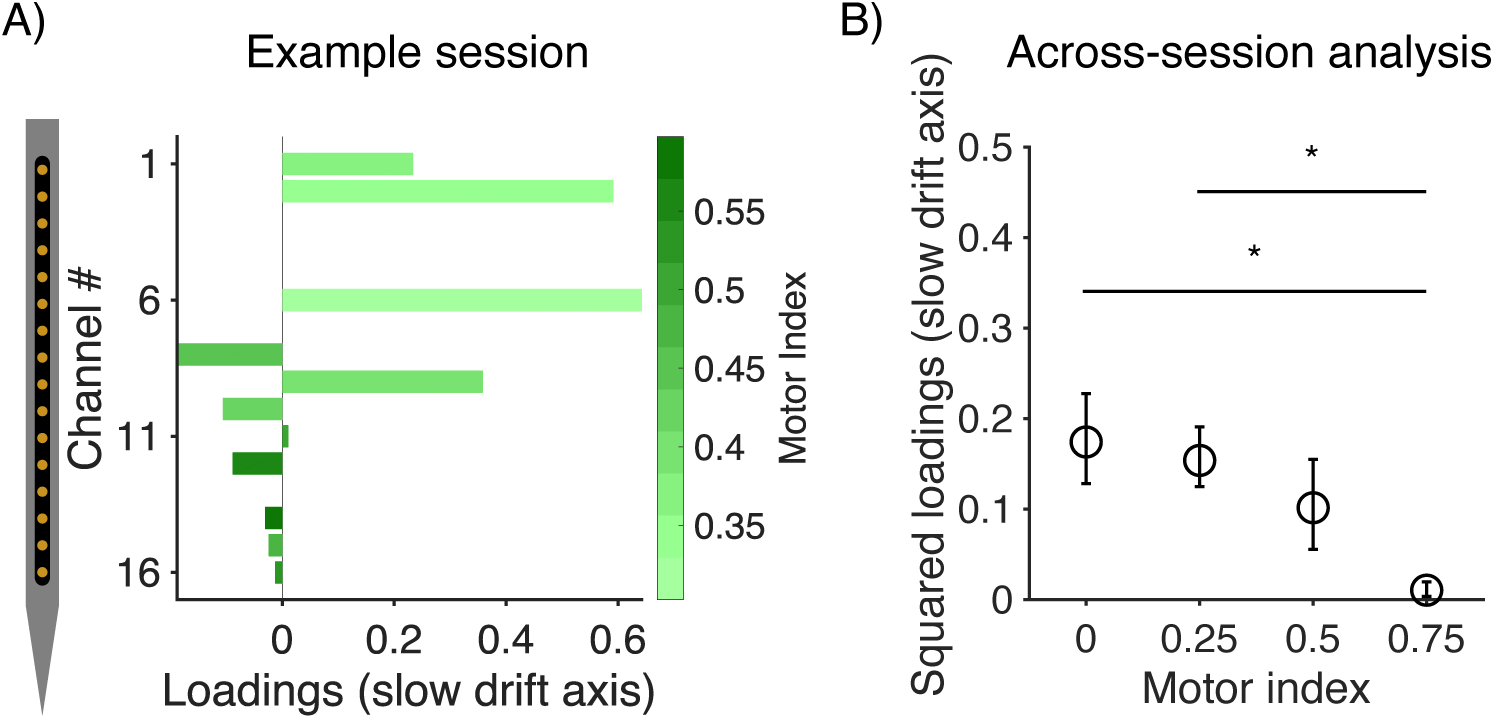
Arousal-related fluctuations are isolated from SC neurons with a higher motor index. (A) Loadings for the slow drift axis for an example session. Motor index is represented by the color bar. Note that loadings were weaker for neurons recorded from channels at the bottom of the probe, which targeted neurons in the deep layers of the SC. These neurons also had higher motor indices, consistent with their role in eye movement generation. (B) Scatter plot for the across session analysis showing squared loadings for the slow drift axis pooled across all recorded SC neurons. The neurons were discretized into four equally spaced bins based on motor index, and permutation tests performed to investigate if neurons with a higher motor index explained less variance in the slow drift axis. The resultant p-values were then corrected for multiple comparisons using Bonferroni adjustment. p < 0.05*, p < 0.01**, p < 0.001***.

To explore if a similar effect was found across sessions, we computed the squared component loading for all recorded SC neurons. This was done to control for the fact that the sign of the correlation between the response of each neuron and the slow drift axis is arbitrary and could be positive or negative (Jolliffe and Cadima, 2016). Next, we discretized the motor indices of all recorded SC neurons into four equally spaced bins. Permutation tests were then performed across the four bins to investigate if neurons with a stronger saccadic response explained less variance in the slow drift axis. After controlling for multiple comparisons (Bonferroni correction), we found that the mean squared component loading for neurons with the highest motor index (M = 0.0105, SD = 0.0237) (i.e. the bin centered on 0.75) was significantly weaker than the mean loading for neurons with lower motor indices (bin centered on 0.25: M = 0.1539, SD = 0.2207; bin centered on 0: M = 0.1743, SD = 0.2429) (Figure 3B). Note that similar results were obtained when motor index was entered as a continuous variable into a multiple regression analysis instead of being discretized into four equally spaced bins (Figure 1 - figure supplement 1). The proportion of variance explained by the model, which also included subject name as a categorical variable to control for intersubject differences, was 0.0350 (F(2, 368) = 6.6748, p = 0.0014). Consistent with our hypothesis, we found that motor index was significantly associated with squared component loadings (t = -3.6154, p = 0.0003) such that SC neurons with a higher motor index explained less variance in the slow drift axis. These results demonstrate that even though arousal-related fluctuations are present in the SC, they are isolated from deep-layers neurons that elicit a strong saccadic response and presumably reside closer to the motor output. As a further test of this hypothesis, we next sought to determine if these same cells explained less variance in projections onto the PFC slow drift axis.

### SC neurons with a higher motor index are less correlated with the PFC slow drift axis

To probe the relationship between deep-layer SC neurons and the PFC slow drift axis, we computed the correlation (Pearson’s product-moment correlation coefficient) between the trial-to-trial residual responses of each SC neuron during the delay period and projections onto the PFC slow drift axis. Consistent with our finding that SC slow drift was significantly correlated with PFC slow drift (Figure 2B), we observed that the trial-to-trial residual responses of some SC neurons were significantly correlated with projections onto the PFC slow drift axis. For example, the residual responses of the neuron shown in Figure 4A, which has a relatively low motor index and is therefore unlikely to reside close to the motor output of the SC, were strongly correlated with projections onto the PFC slow drift axis (r^2^ = 0.5346, p < 0.001). However, other neurons showed no correlation such as that in Figure 4B. The residual responses of this neuron, which has a relatively high motor index and presumably resides closer to the motor output stage, were not significantly correlated with projections onto the PFC slow drift axis (r^2^ = 0.0010, p = 0.2650).

**Figure 4.**
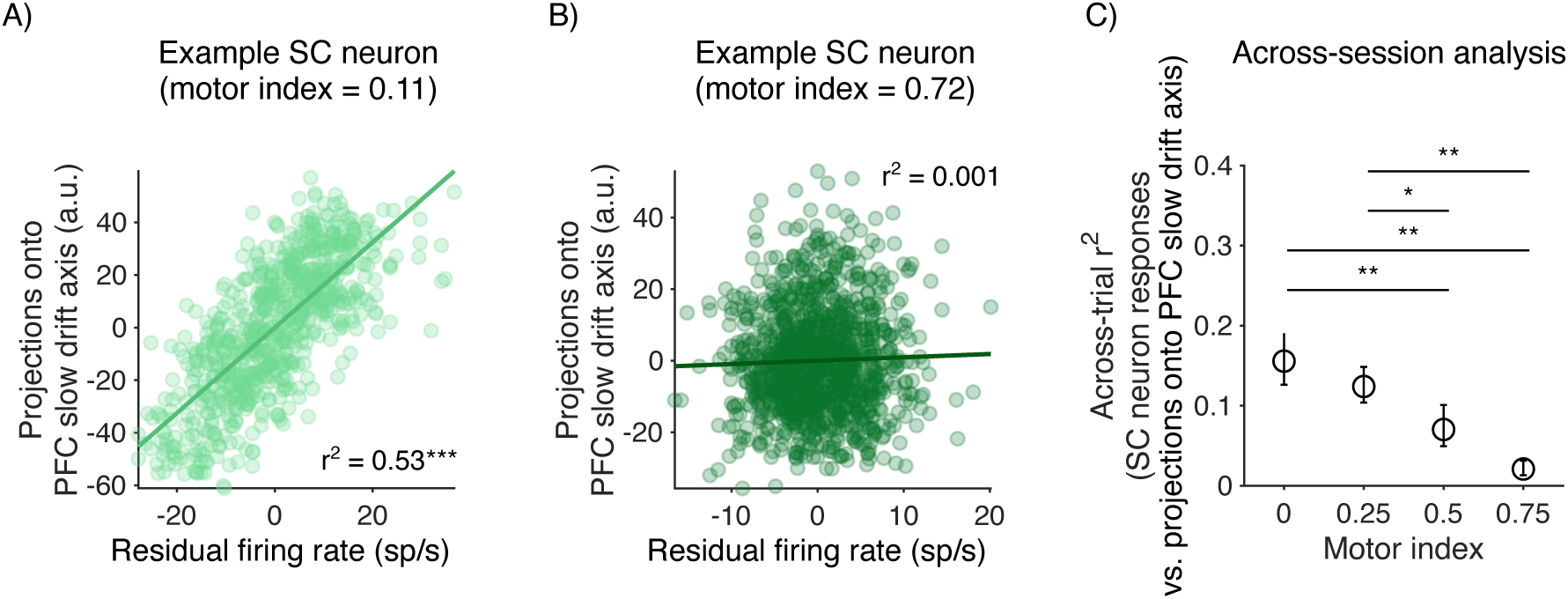
SC neurons with a higher motor index are less correlated with the PFC slow drift axis. (A) Example SC neuron with a relatively low motor index that exhibited a significant correlation between its residual spiking response and projections onto the PFC slow drift axis (r^2^ = 0.5346, p < 0.001). (B) Example SC neuron with a relatively high motor index that did not exhibit a correlation between its residual spiking response and projections onto the PFC slow drift axis (r^2^ = 0.001, p = 0.2650). (C) Scatter plot for the across session analysis showing the mean variance explained between residual spiking responses and projections onto the PFC slow drift axis for all recorded SC neurons. The neurons were discretized into four equally spaced bins based on motor index, and permutation tests performed to investigate if the variance explained between residual SC spiking responses and projections onto the PFC slow drift was lower for neurons with a higher motor index. The resultant p-values were then corrected for multiple comparisons using Bonferroni adjustment. p < 0.05*, p < 0.01**, p < 0.001***.

To explore if a similar effect was found across sessions, we again pooled the motor indices of all recorded SC neurons and discretized them into four equally spaced bins. We performed permutation tests across the bins to investigate if the mean variance explained between residual SC neuron responses and projections onto the PFC slow drift axis was lower for neurons with a higher motor index. After controlling for multiple comparisons, we found that the mean variance explained was significantly lower for neurons with the highest motor index SC (M = 0.0209, SD = 0.0236) (i.e. the bin centered on 0.75) than it was for neurons with lower motor indices (bin centered on 0.25: M = 0.1240, SD = 0.1468; bin centered on 0: M = 0.1557, SD = 0.1591) (Figure 4C). Furthermore, the mean variance explained for neurons with an intermediate motor index (i.e. the bin centered on 0.5) (M = 0.0703, SD = 0.1095) was significantly lower than it was for neurons with the lowest motor indices (i.e. the bins centered on 0.25 and 0). Once again, similar results were obtained when the motor index was entered as a continuous variable into a multiple regression model instead of being discretized into bins (Figure 1 - figure supplement 1). The proportion of variance explained by the model, after controlling intersubject differences, was 0.1777 (F(2, 368) = 39.7739; p < 0.001). Motor index was significantly associated with the strength of the relationship between single SC neurons responses and projections onto the PFC slow drift axis (t = -2.7902; p = 0.0055) such that it was weaker for neurons with a higher motor index. These findings provide additional support for the hypothesis that arousal-related fluctuations are isolated from neurons in the deep layers of the SC. They are more characteristic of cells residing at the early to mid stages of the visuomotor transformation, which have been implicated in autonomic processes such as controlling the size of the pupil (Wang et al., 2012; Wang and Munoz, 2018). This prompted us to ask if a relationship exists between SC neurons responses and pupil size, and whether this relationship is mediated by motor index.

### The relationship between SC neuron responses and pupil size is modulated by motor index

Previous work has shown that intermediate-layer SC neurons control the size of the pupil via: 1) direct projections to the Edinger-Westphal nucleus; and 2) indirect projections to the mesencephalic cuneiform nucleus (Joshi and Gold, 2020). Thus, we hypothesized that neurons with a lower motor index would explain more variance in trial-to-trial pupil size than neurons in deep layers. To test this, we computed the correlation (Pearson’s product-moment correlation coefficient) between the spiking responses of each SC neuron and pupil size, both of which were computed during the first 200ms of the delay period. In several sessions, we found that the strength of this relationship was inversely related to motor index. For example, the spiking response of the neuron shown in Figure 5A, which has a relatively low motor index and presumably resides at the mid stage of the visuomotor transformation, was strongly correlated with pupil size (r^2^ = 0.2121, p < 0.001). In contrast, the spiking responses of the neuron shown in Figure 5B, which has a relatively high motor index and presumably resides closer to the motor output stage, was not significantly correlated with changes in pupil size over the course of a recording session (r^2^ = 0.001, p = 0.7534).

**Figure 5.**
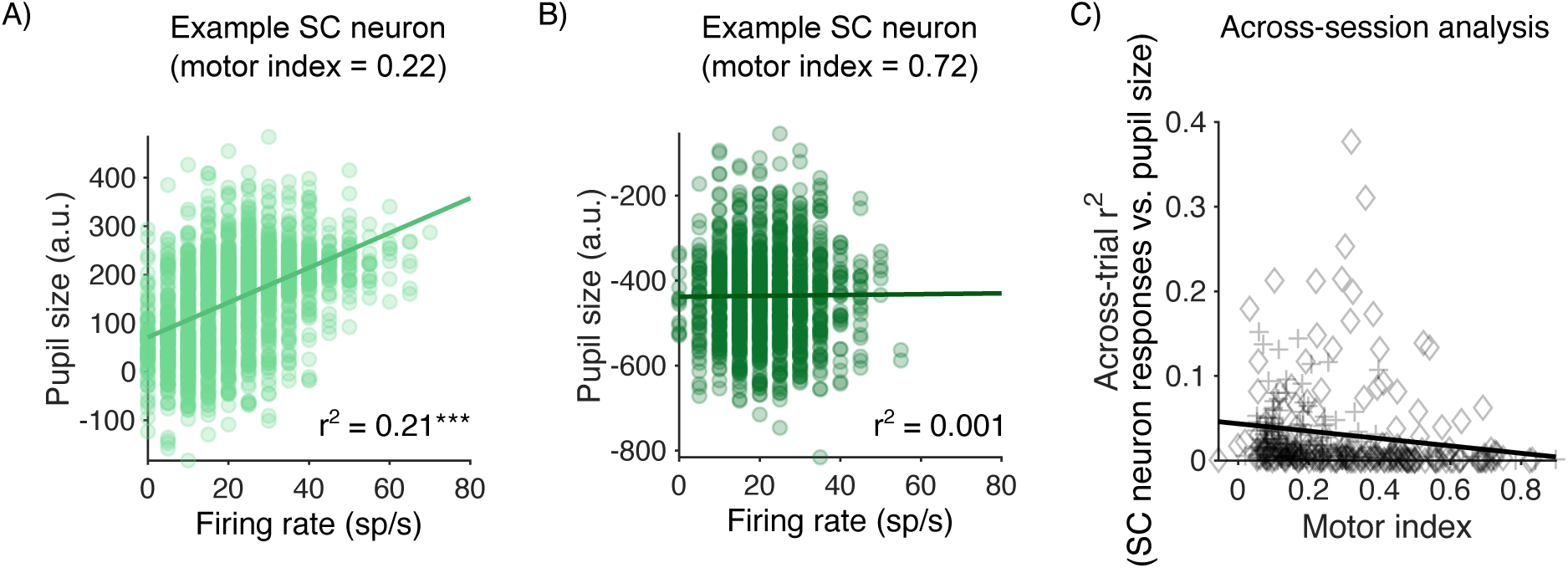
The relationship between SC neuron responses and pupil size is modulated by motor index. (A) Example SC neuron with a relatively low motor index that exhibited a significant correlation between its spiking response and pupil size. (B) Example SC neuron with a relatively high motor index that did not exhibit a correlation between its spiking response and pupil size (r^2^ = 0.1601, p < 0.001). (C) Scatter plot showing that the strength of the relationship between individual SC neuron responses and pupil size is inversely related to motor index (r^2^ = 0.0309, p = 0.0031). p < 0.05*, p < 0.01**, p < 0.001***.

To determine if a similar effect was found across sessions, we performed a multiple regression analysis. In this analysis, we sought to determine if motor index predicted the strength of the relationship between SC neuron responses during the delay period and pupil size, as measured using the coefficient of determination (r^2^). After controlling for intersubject differences, results showed that the proportion of variance explained by the model was 0.0309 (F(2, 368) = 5.8717; p = 0.0031). Motor index was significantly associated with the strength of the relationship between SC neuron responses and pupil size (t = -3.2593; p = 0.0012) (Figure 5C). More specifically, we found a stronger relationship between neuronal response and pupil size for neurons with a lower motor index, consistent with the proposed role of intermediate-layer SC neurons in controlling the size of the pupil (Joshi and Gold, 2020).

We also considered whether shifts in SC neural activity could be driven directly by changes in luminance induced by changes in the size of the pupil. If such luminance sensitivity existed in SC among the visually selective neurons, it could explain the slow fluctuations in their activity correlating with slow fluctuations in the size of the pupil. Neurons in the SC are tuned for transient changes in stimulus contrast (Chen and Hafed, 2018), though we are not aware of studies examining how slow shifts in overall luminance affect SC responses. Nonetheless, to ask whether this might be the case, we performed two analyses. First, we analyzed the pupil size in a different period of time from the neural activity, the baseline period before target presentation. Although there was only a fixation point on the screen at that time and no target to induce a transient visual response to luminance, we found similar results relating SC spiking responses to pupil size (Figure 5 - figure supplement 1). Second, we compared slow drift measures computed on spiking responses during different epochs, and found they were highly correlated (Figure 2 – figure supplement 1). Because slow drift is computed on an average spiking across many trials, this makes it unlikely that transient luminance-induced responses could lead to changes in these fluctuations. Given that slow drift is found in traditionally defined visual areas (e.g., area V4) and in regions that show mixed selectivity for multiple task variables (e.g., PFC) (Cowley et al., 2020), it seems unlikely that slow drift is caused by luminance fluctuations alone and more likely that it reflects global changes in arousal. At the same time, these arousal-related fluctuations covary with changes in pupil size (Johnston et al., 2022a), which could modulate the amount of light entering the eye from the display. This might affect visual neurons but not motor neurons due to their lack of visual sensitivity. Because SC neurons exist on a continuum, with visual responses decreasing and motor responses increasing from the intermediate to deep layers (Massot et al., 2019; Heusser et al., 2022) and no clear categorical boundary for motor-only neurons, any readout strategy would still need to avoid corruption of the motor output by slow drift, even if it were caused by changes in the amount of light entering the eye.

Taken together, the findings outlined above suggest that arousal-related signals are kept separate from neurons in the deep layers of the SC. In addition to being more weakly correlated with pupil size, the spiking responses of these neurons did not exhibit large fluctuations over time (Figure 2), and when considering the neuronal population as a whole, explained less variance in the slow drift axis when it was computed using population activity in the SC (Figure 3) and PFC (Figure 4). Although these results point to a segregation of arousal- and eye movement-related signals in the SC, research has shown that all three layers receive input from the LC: a brain region that has long been associated with controlling global arousal levels (Benavidez et al., 2021). Thus, an additional mechanism may have evolved to ensure that arousal-related signals are kept separate from those used in the encoding of eye position. In primary motor cortex, signals related to movement preparation reside in an orthogonal axis in population activity space to those that are used for movement initiation so that they do not lead to premature muscle contractions (Kaufman et al., 2014; Elsayed et al., 2016). Next, we explored if a similar mechanism exists in the SC to prevent arousal-related signals interfering with the motor output.

### Arousal-related signals in the SC reside in an orthogonal subspace to saccade-related signals

To investigate if arousal-related signals in the SC reside in an orthogonal subspace to those that are used to encode eye position, we computed a “pupil size axis” for each session. We discretized trial-to-trial pupil size into eight equally spaced bins and, for each neuron, computed the mean spiking response per bin (Figure 6A, left). We then used PCA to find the axis in population activity space that explained the most variance in the data attributable to the pupil. The loadings (Figure 6A, right) represent the correlation between the response of each neuron across bins and the first principal component meaning that neurons exhibiting a stronger relationship between firing rate and pupil size (e.g. Figure 6A, Channel 5) explained more variance in the pupil size axis. Our goal was to compare projections onto the pupil size axis to projections onto a “saccade tuning axis”. To compute this axis, tuning curves were generated for each SC neuron by taking the mean spiking response during the saccade epoch for each target angle (Figure 6B, right). PCA was then applied to the tuning curves, which yielded a vector of loadings for the first principal component (Figure 6B, left). In this case, neurons that exhibited greater selectivity for a particular target angle explained more variance in the saccade tuning axis (e.g. Figure 6B, Channel 11).

**Figure 6.**
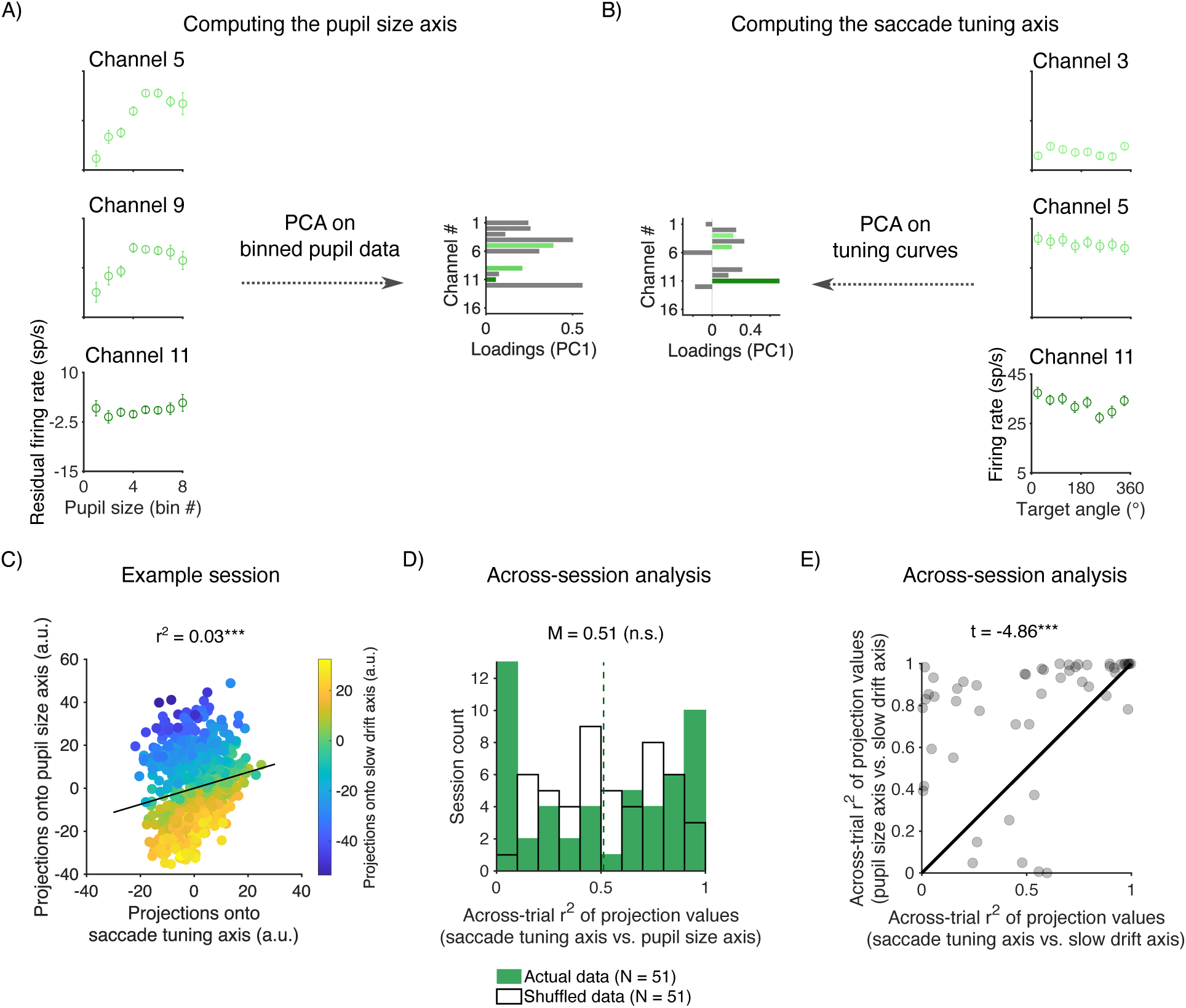
Arousal-related signals in the SC reside in an orthogonal subspace to saccade-related signals. (A) Computing the pupil size axis for an example session. Three example SC neurons from the same recording session (left). Each data point represents the mean spiking response after trial-to-trial pupil size had been discretized into eight equally spaced bins. Note that pupil size increases with bin number such that bin 1 contains trials in which the pupil was most constricted, whereas bin 8 contains trials in which the pupil was most dilated When PCA was applied to the data it yielded a vector of loadings for the first principal component (right). (B) Computing the saccade tuning axis for the same session shown in (A). Three example SC neurons (right). Each point represents the mean spiking response during the saccade epoch (25ms before to 25ms after the onset of the saccade) for each target angle. When PCA was applied to the data it yielded a vector of loadings for the first principal component (left). (C) A scatter plot showing the relationship between projections onto the pupil size axis and the saccade tuning axis for the same example session shown in (A) and (B). The color scale represents projections onto the slow drift axis. Note that the same data (i.e. residual SC responses during the delay period) were projected onto the three axes. (D) Histogram showing the proportion of variance explained between projections onto the pupil size axis and projections onto the saccade tuning axis. A p-value was computed by comparing the actual distribution of values to a null distribution (see Methods). (E) Scatter plot comparing the variance explained between projections onto the pupil size axis and the slow drift axis, and projections onto the saccade tuning axis and the slow drift axis. A p-value was computed using a paired-sample t-test. p < 0.05*, p < 0.01**, p < 0.001***.

We first investigated if the pupil size axis and the saccade tuning axis reside in orthogonal subspaces. To ensure that fair comparisons could be made, the same data (i.e. trial-to-trial residual responses during the delay period) were projected onto each of the axes. We then computed a correlation (Pearson’s product-moment correlation coefficient) between the projection values for each session, and used the coefficient of determination (r^2^) to assess the variance explained. If arousal-related signals in the SC reside in an orthogonal subspace to those that are used for saccade generation, one would expect projections onto the pupil size axis to be weakly correlated with projections onto the saccade tuning axis. This was found to be the case in several sessions such as that shown in Figure 6C (r^2^ = 0.0307, p < 0.001). Furthermore, when the data were pooled across sessions, we found that the mean variance explained between the projection values was 0.5142 (Figure 6D). By comparing the actual distribution of r^2^ values to a null distribution that was generated by correlating projections onto the pupil size axis with projections onto an axis that was randomly oriented in population activity space, we were able to show that the mean variance explained was not significantly greater than what would be expected for data projected onto a random axis (p = 0.5192).

Finally, as an additional test of our hypothesis, we investigated if the pupil size axis was more aligned to the slow drift axis than the saccade tuning axis. To do this, we projected trial-to-trial residual responses during the delay period onto the pupil size axis and the saccade tuning axis. For each session, we then correlated (Pearson’s product-moment correlation coefficient) these projection values with projections onto the slow drift axis. Across sessions, we found that the mean variance explained between projections onto the pupil size axis and the slow drift axis (M = 0.7780, SD = 0.3025) was significantly greater than the mean variance explained between projections onto the saccade tuning axis (M = 0.5319, SD = 0.3407) (t = -4.8578, p < 0.001).

## Discussion

The main aim of this study was to identify how global fluctuations in arousal affect subcortical areas like the SC, and how such fluctuations interact with the motor functions of that area. In addition to representing a highly conserved brain structure in vertebrates, evidence suggests that the SC also has a number of divergent properties that have been linked to higher-order cognitive processes such as spatial attention (Knudsen, 2011; Krauzlis et al., 2013; Clark et al., 2015; Basso et al., 2021) and perceptual decision-making (Ratcliff et al., 2003; Horwitz et al., 2004; Crapse et al., 2018; Jun et al., 2021). In a similar way to how cortical motor areas separate out preparatory signals (Kaufman et al., 2014; Elsayed et al., 2016) and signals from the opposing arm (Ames and Churchland, 2019) when performing a reach movement, oculomotor structures such as the SC must be able to disentangle signals related to cognition and brain state from those used for eye movement encoding.

Consistent with what has been observed in the cortex (Cowley et al., 2020), we found that the activity of some SC neurons fluctuated over time, and by reducing the dimensionality of the data we were able to isolate a neuronal signature of arousal that was correlated with pupil size and simultaneously recorded PFC data. In the context of the SC, this motivated us to ask how the precise coordination of saccades is maintained despite the presence of ongoing fluctuations in brain state. Another internal cognitive state that has been studied in the context of the visual and motor role of the SC is spatial attention. Research using single-unit recordings in monkeys (Ignashchenkova et al., 2004) and neuroimaging in humans (Katyal et al., 2010) has shown that the effects of spatial attention on SC activity are layer-specific and more characteristic of neurons in the superficial and intermediate layers. At a population level, attention has been shown to increase the efficacy of communication from MT to SC in a similar manner for visual and motor SC populations (Srinath et al., 2021) and reshape the visual information provided by MT to SC (Ruff and Cohen, 2019). Overall, both spatial attention and global arousal share the property of having an influence on the eye movement system while at the same time needing to avoid directly producing unwanted eye movements. Here, we propose that arousal-related signals in the SC are segregated from the motor planning function of SC to prevent unwanted changes in eye position. To test this hypothesis, we utilized linear array recordings to simultaneously monitor the activity of SC neurons with distinct neurophysiological response properties (Massot et al., 2019; Ayar et al., 2023). Across sessions, we found that neurons exhibiting a stronger saccadic response, which largely reside in the deep layers of the SC closer to the motor output (Basso and May, 2017), were less connected to SC slow drift measured in the population response. Furthermore, by correlating the response of these cells with a slow drift axis that was generated using simultaneously recorded PFC data, we were able verify the global nature of this signal. These findings are consistent with neuroimaging studies showing that changes in brain state lead to coordinated fluctuations of activity across the brain (Liu et al., 2018; Turchi et al., 2018; Setzer et al., 2022), and research that recorded simultaneously from neurons in cortical areas and subcortical areas such as the LC (Joshi and Gold, 2022; Collins et al., 2023). Furthermore, our results suggest that arousal-related fluctuations are more characteristic of neurons residing in the intermediate layers of the SC. In support of this, several studies have pointed to a link between the activity of these cells and non-invasive markers of arousal such as pupil size (Wang et al., 2012, 2014; Joshi et al., 2016; Wang and Munoz, 2021).

To further study the role of intermediate-layer SC neurons in controlling the size of the pupil, we investigated if the strength of the relationship between trial-to-trial changes in pupil size and SC spiking responses was mediated by motor index. Consistent with the proposed link between intermediate-layer neurons and pupil size, we found that cells exhibiting a weaker saccadic response explained more variance in pupil size than neurons with a stronger saccadic response. These results provide further support for the notion that arousal-related signals are kept separate from signals used to encode eye position in the SC. This provides a potential mechanism through which signals related to cognition and brain state can exist in oculomotor structures without affecting high precision fixation and eye movements.

In addition to being kept separate at the level of different layers in the SC, signals related to arousal and eye position encoding may also occupy different dimensions in population activity space. The term “subspace” is often used when referring to the fact that the activity of simultaneously recorded neurons is correlated (Cohen and Kohn, 2011) and occupies fewer dimensions than the number of neurons in the population (Ebitz and Hayden, 2021; Rust and Cohen, 2022). The concept of a subspace has been used to identify how populations of neurons segregate different types of activity at the population level, rather than simply in different neurons. Non-overlapping subspaces can be used to encode different types of sensory information (Mante et al., 2013; Aoi et al., 2020; Ebitz et al., 2020), facilitate communication between distinct neural circuits (Semedo et al., 2019, 2022), and isolate preparatory signals from movement commands (Churchland and Shenoy, 2024). For example, when recording from neurons in the motor cortex while monkeys performed a delayed-reach task, (Kaufman et al., 2014) found that preparatory signals reside in an orthogonal subspace to those used to encode reach direction. In the SC, (Ayar et al., 2023) found that signals related to sensation and action occupy separate subspaces and depend upon the cognitive demands of the task, consistent with previous research showing that the temporal structure of the SC population response differs during visual and motor epochs (Jagadisan and Gandhi, 2022). To explore if a similar mechanism exists in the SC, we projected trial-to-trial responses onto a pupil size axis and a saccade encoding axis, and found that the variance explained was not significantly greater than what would be expected for data projected onto a random axis. Furthermore, projections onto the pupil size axis were more aligned to projections onto the slow drift axis than projections onto the saccade encoding axis. Taken together, these findings suggest that arousal-related signals reside in an orthogonal subspace to those used to encode eye movement plans. In a similar manner to how preparatory signals in the motor cortex do not lead to premature muscle contractions (Kaufman et al., 2014; Elsayed et al., 2016), this might explain why arousal-related signals in the SC do not lead to unwanted changes in eye position.

In summary, by simultaneously recording from populations of neurons in the SC and the PFC, we were able to observe a fluctuation of activity that was present in both areas and correlated with non-invasive markers of arousal such as pupil size. The presence of arousal-related signals in the SC is consistent with its role in cognitive processes such as spatial attention and decision-making. However, it also raises the question of how motor areas can disentangle these higher-order signals from those used for movement encoding. In addition to being kept separate at the level of different layers in the SC, our results demonstrate that signals related to arousal and eye movement planning occupy distinct dimensions in population activity space. Further research is needed to determine whether this represents a similar mechanism to that which is used in the motor cortex to separate preparatory signals from movement commands (Churchland and Shenoy, 2024). If so, it could provide an explanation for how cognitive signals can be intermixed and impact activity in populations of SC neurons without affecting precision in fixation and saccadic eye movements.

## Methods

### Subjects

Two adult rhesus macaque monkeys (Macaca mulatta) were used in this study (Monkey Do = 28 sessions, Monkey Wa = 34 sessions). Surgical procedures to chronically implant a titanium head post (to immobilize the subjects’ heads during experiments) and microelectrode arrays were conducted in aseptic conditions under isoflurane anesthesia. Recording chambers situated along the midline at an angle of 30° from vertical and aimed at a location 8mm above interaural zero were also implanted for SC recordings. Opiate analgesics were used to minimize pain and discomfort during the perioperative period. Experimental procedures were approved by the Institutional Animal Care and Use Committee of Carnegie Mellon University and were performed in accordance with the United States National Research Council’s Guide for the Care and Use of Laboratory Animals.

### Superior colliculus (SC) recordings

The spiking responses of neuronal populations in the SC were recorded using a 16-channel linear microelectrode array (V-Probe, Plexon, Dallas, TX). The total length of the linear array was 115mm. The contacts were spaced 150μm apart and spanned a distance of 2.25mm. Linear arrays were lowered into the SC through a guide tube using a custom-designed mechanical microdrive (Laboratory for Sensorimotor Research, Bethesda, MD), which was inserted through a plastic grid with 1mm spacing.

### Prefrontal cortex (PFC) recordings

The spiking responses of neuronal populations in the PFC (Monkey Do = left hemisphere, Monkey Wa = right hemisphere) were recorded using 96-channel “Utah” arrays (Blackrock Microsystems). The arrays comprised a 10x10 grid of silicon microelectrodes (1mm in length) that were spaced 400μm apart, implanted on the pre-arcuate gyrus (8Ar).

### Waveform classification

Signals from the linear probes (SC) and Utah arrays (PFC) were amplified and band-pass filtered (0.3–7500Hz) by a Grapevine system (Ripple). Waveform segments crossing a threshold (set as a multiple of the root mean square noise on each channel) were digitized (30KHz) and stored for offline processing, which involved waveforms being automatically sorted as spikes or noise by a neural network classifier developed in our laboratory (code available at https://github.com/smithlabvision/spikesort/nasnet). Although this method cannot distinguish between multiple neurons recorded from the same channel, it provides fast and accurate classifications that are highly similar to those of a human spike-sorter (Issar et al., 2020). All waveforms recorded from a single channel that remained after the neural network classifier had been applied were considered to be from a single unit, which we refer to as a “neuron”.

### Exclusion criteria

Across both monkeys, we recorded from 992 sites in SC and 5952 sites in PFC. This included all channels from the 16-channel linear microelectrode array and all channels from the 96-channel utah array across all sessions (SC: Monkey Do = 448, Monkey Wa = 544; PFC: Monkey Do = 2688, Money Wa = 3264). We applied a rigorous set of exclusion criteria designed to eliminate neurons with noisy or unstable waveforms, those with low firing rates, and neurons that did not have an evoked response during either the visual or motor epoch of the MGS

To identify candidate neurons that were mostly classified as containing noise waveforms by the neural network classifier, and therefore only comprised a small number of waveforms, we removed neurons that had a mean firing rate less than 2sp/s across the entire session (Smith and Sommer, 2013). This was calculated by taking the total number of spikes in the recording session from the beginning (before the task was started) to the end (after the last trial of the day), and dividing by the total time (including periods of task performance and periods outside of the trial). In addition to removing neurons that had a low mean firing rate across an entire session, we also removed SC neurons that did not exhibit elevated firing rates during the visual and motor epochs of the MGS tasks i.e. following the presentation of the target stimulus and during the onset of a saccade. This was quantified by computing a visual and motor index for each SC neuron (see below). Neurons were removed if the value of both of these indices was less than 0.05, indicating a lack of response during both epochs. To rule out the possibility that slow drift arose due to recording instabilities (e.g. the distance between the neuron and the microelectrode(s) changing over time), we only included neurons with stable waveform shapes throughout a session. Neurons were removed if they had a percent waveform variance greater than 10% (Cowley et al., 2020). We also removed neurons that exhibited abrupt changes in activity during a recording session. This was quantified by dividing each session into 20 non-overlapping time bins and computing the mean spike count per bin (Cowley et al., 2020). Neurons were removed if the change in activity between any two consecutive time bins was greater than 25% of the maximum activity of that neuron. Adopting these exclusion criteria ensured that slow drift did not arise due to recording instabilities. We also note that if drift were to occur due to recording instabilities, it would not be expected to affect neurons from the same penetration differentially based on their motor index (e.g., Figure 3A).

In order to obtain a precise estimate of the slow drift axis, and after controlling for low firing rate and recording instabilities, we removed sessions that contained less than two SC or PFC neurons (Monkey Do = 3 sessions, Monkey Wa = 5 sessions). Furthermore, we removed sessions if they had a duration lasting less than 45 minutes (Monkey Do = 0 sessions, Monkey Wa = 3 sessions). Adopting these additional criteria ensured that there were enough variables and observations to perform PCA and accurately compute the slow drift axis. Using all of this criteria, the total number of sessions included in the analysis for Monkey Do and Monkey Wa was 25 sessions (SC = 190 neurons, PFC = 795) and 26 sessions (SC = 181, PFC = 2110), respectively.

### Visual stimuli

Visual stimuli were generated using a combination of custom software written in MATLAB (The MathWorks) and Psychophysics Toolbox extensions (Brainard, 1997; Pelli, 1997; Kleiner et al., 2007). They were displayed on a CRT monitor (resolution = 1024 X 768 pixels; refresh rate = 100Hz), which was viewed at a distance of 57cm and gamma-corrected to linearize the relationship between input voltage and output luminance using a photometer and look-up-tables.

### Memory-guided saccade task

Subjects fixated a central point (diameter = 0.18°) on the monitor to initiate a trial (Figure 1A). After fixating within a circular window (diameter: Monkey Do = 2.38°, Monkey Wa = 1.10°) for 200ms, a target stimulus (diameter = 0.55°) was presented for 200ms at one of eight angles separated by 45°. The eccentricity of the target stimulus was determined on a session-by-session basis by applying low-amplitude microstimulation to contacts along the linear microelectrode array to evoke saccadic eye movements. Target eccentricity was then set such that it matched the endpoints of evoked saccades (M = 4.52°, SD = 2.89°). After the target stimulus had been presented, subjects were required to maintain fixation for a delay period. The duration of the delay period chosen at random on each trial from a uniform distribution spanning 600-1100ms. If steady fixation was maintained throughout the delay period, the central point was extinguished prompting the subjects to make a saccade to the remembered target location. The subjects had 500ms to initiate a saccade. To receive a liquid reward, the subjects’ gaze had to be maintained within a circular window centered on the remembered target location (diameter: Monkey Do = 2.93°, Monkey Wa = 2.20° for the first 12 sessions and 3.01° for the remaining 22 sessions) for 100ms. In a subset of sessions, the target was briefly reilluminated, after the fixation point was extinguished and the saccade had been initiated, to aid in saccade completion.

### Eye tracking

Eye position and pupil diameter were recorded monocularly at a rate of 1000Hz using an infrared eye tracker (EyeLink 1000, SR Research).

### Microsaccade detection

Microsaccades were detected during the delay period and were defined as eye movements that exceeded a velocity threshold of 6 times the standard deviation of the median velocity for at least 6ms (Engbert and Kliegl, 2003) (Figure 2 - figure supplement 2). To prevent overshoots (i.e. fast, oppositely directed gaze shifts that occur shortly after a microsaccade) being treated as separate eye movements, we merged microsaccades that occurred within 50ms of each other (McCamy et al., 2015). Finally, we assessed the validity of our microsaccade detection method by computing the correlation (Pearson’s product-moment correlation coefficient) between the amplitude and peak velocity of detected microsaccades for each session. The mean correlation between these two variables across sessions was 0.7601 (SD = 0.0678) indicating that our detection algorithm was robust, as microsaccades fell on the main sequence (Zuber et al., 1965).

### Characterizing the response properties of SC neurons

To characterize the neurophysiological response properties of individual SC neurons, we first visualized SC responses by computing PSTHs aligned to the onset of the target stimulus and the onset of the saccade to the remembered target location, respectively (Figure 1B). Firing rates were computed in 1ms bins, averaged across trials and smoothed using a Gaussian function (σ = 5ms).

To further quantify the strength of the saccadic response to the remembered target location, a motor index was computed for each neuron as follows:

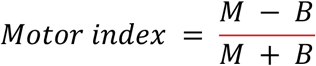

Where *M* represents the mean response during a peri-saccade epoch from 25ms before to 25ms after the onset of the saccade, and *B* represents the mean response during a baseline period defined as 100ms before the onset of the target stimulus. Higher motor index values are indicative of a stronger saccadic response relative to baseline, whereas lower values represent a weaker saccadic response relative to baseline. Note that we also computed a visual index for each neuron, which was only used for the purpose of excluding neurons that did not exhibit an evoked response during the MGS task (see above). The visual index was computed in the following manner:

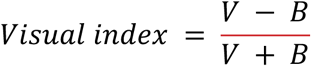

Where *V* represents the mean response during a peri-stimulus epoch spanning 25ms to 125ms after the onset of the target stimulus, and *B* represents the mean response during a baseline period defined as 100ms before the onset of the target stimulus.

## Generating shuffled distributions

### Variance explained by SC slow drift axis across sessions

To determine if the variance explained by the SC slow drift axis was significantly greater than chance across sessions, the actual distribution of proportion variance explained values was compared to a shuffled distribution (Figure 2A, top right). To generate this distribution, PCA was performed on binned spike counts that had first been shuffled across trials (1000 iterations). We then computed the mean variance explained by the first principal component across iterations, which yielded a single shuffled value for each session. Finally, a p-value was computed by calculating the proportion of values in the shuffled distribution that were greater than or equal to the actual variance explained across sessions.

### Relationship between SC slow drift axis and the PFC slow drift

To assess the significance of the relationship between projections onto the SC slow drift axis and the PFC slow drift axis (Figure 2B), we randomly shuffled the two variables across sessions and re-computed the correlations (1000 iterations). Differences in session length were controlled for by truncating longer sessions such that they comprised the same number of data points as the shorter session. A p-value was then computed by calculating the proportion of values in the shuffled distribution that were greater than or equal to the actual variance explained across sessions. Note that an identical method was used to determine if projections onto the SC slow drift axis explained a significant amount of the variance in pupil size (Figure 2C) and microsaccade rate (Figure 2 - figure supplement 2).

### Orthogonality of the pupil size axis and the saccade tuning axis

To determine if the pupil size axis and the saccade tuning axis reside in orthogonal subspaces, we computed the correlation between projections onto each axis and then compared the actual distribution of r^2^ values across sessions to a shuffled distribution (Figure 6D). To generate this distribution, projections onto the pupil size axis were correlated with projections onto an axis that was randomly orientated in population activity space (1000 iterations). The latter was generated in a similar manner to the saccade tuning axis, the only difference being that tuning curves were computed for target angles that had been shuffled across trials. We then computed the mean variance explained across iterations, which yielded a single shuffled value for each session. Finally, a p-value was computed by calculating the proportion of values in the shuffled distribution that were greater than or equal to the actual variance explained across sessions.

## Acknowledgments

The authors would like to thank Samantha Schmitt for assistance with surgery and data collection, Karen McCracken for assistance with animal care, and Dr. Katerina Acar for advice about task design and assistance with the mapping of the recording chamber. We would also like to thank Dr. Siddhartha Joshi and Dr. Neeraj Gandhi for providing expert guidance about chamber placement. M. S. was supported by NIH R01 EY029250 and NIH R01MH128393.

---

**Figure 1 - figure supplement 1.**
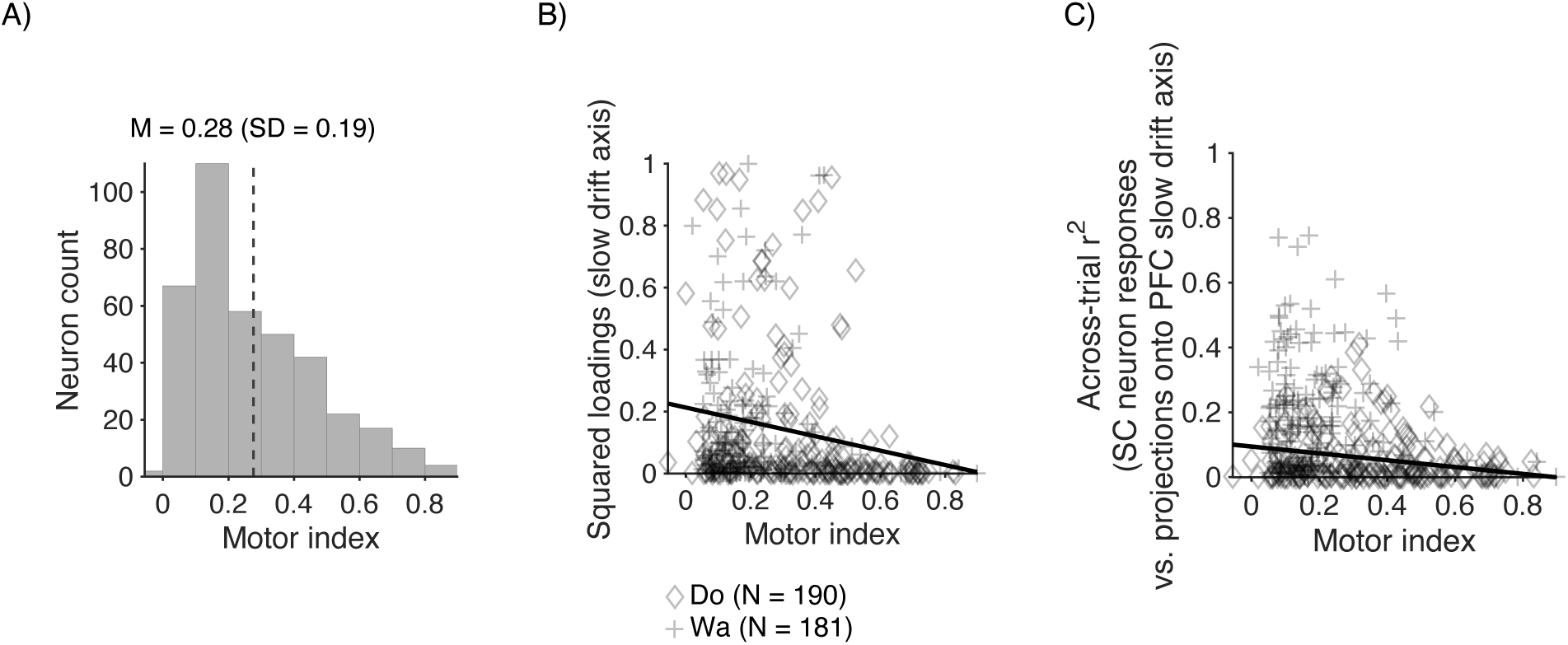
(A) Histogram showing the distribution of motor index values computed across sessions for all recorded SC neurons. The mean value across sessions is denoted by the dashed line. (B) Scatter plot showing the relationship between motor index and squared component loadings for the SC slow drift axis. The line of best fit was computed using multiple regression. Motor index was entered as a continuous predictor variable along with subject name to control for differences between subjects. As described in the Results section, the model explained a significant amount of the variance in the data. A negative correlation was also found between motor index and squared loadings. (C) Scatter plot showing the relationship between motor index and the strength of the relationship between single SC neurons responses and projections onto the PFC slow drift axis, as measured by computing r^2^. As in (B), the line of best fit was computed using multiple regression. Results showed that the model explained a significant amount of the variance in the data. Furthermore, a negative correlation was found between motor index and the r^2^ between single neuron SC responses and projections onto the PFC slow drift axis.

**Figure 2 - figure supplement 1.**
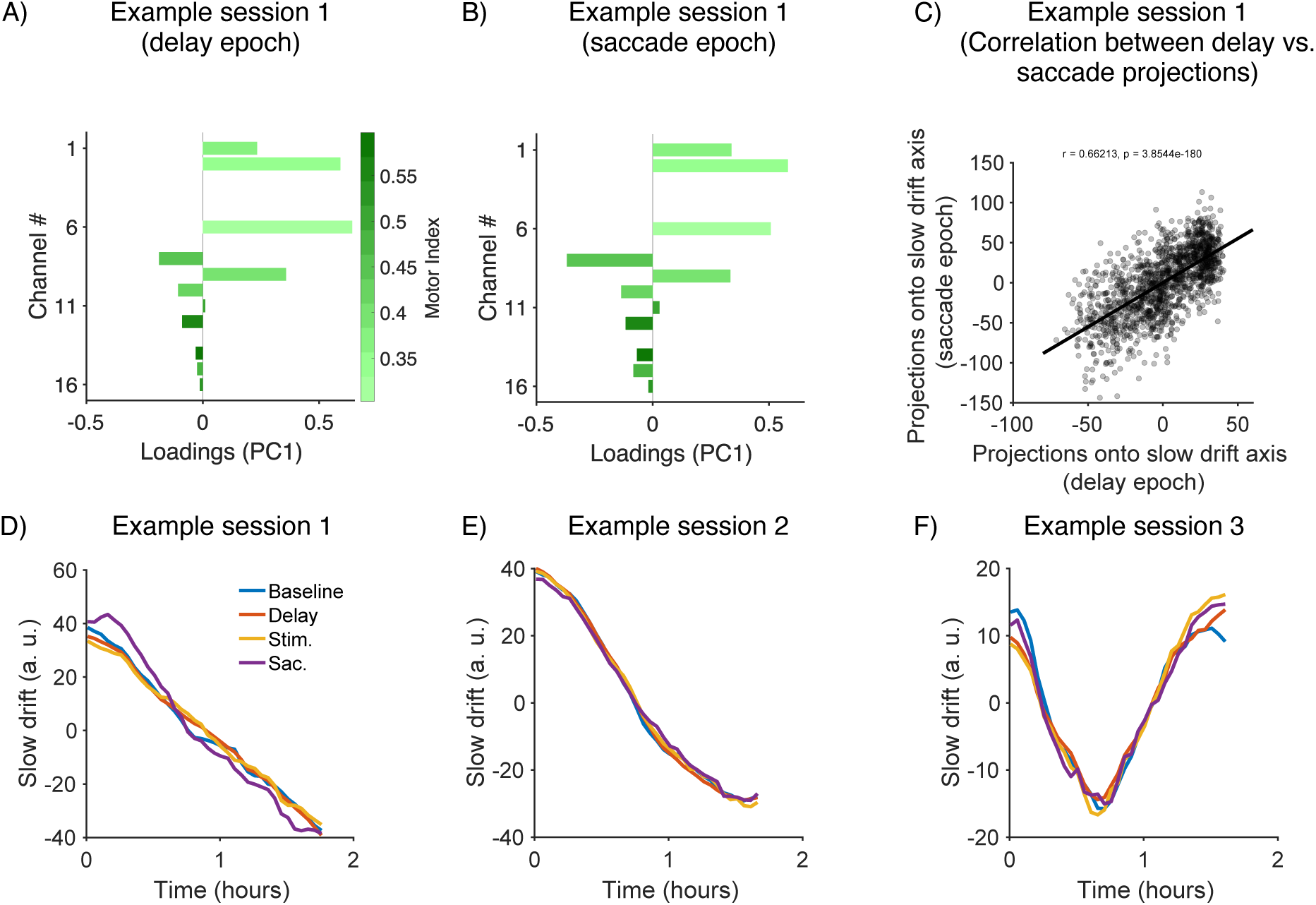
(A) Loadings for the SC slow drift axis for an example session when it was computed using spiking responses during the delay period (same session as Figure 3A in main manuscript). (B) Loadings for the SC slow drift axis from the same session when it was computed using spiking responses during the saccade epoch. (C) Scatter plot showing the correlation between projections onto the slow drift axis for the same session when the slow drift axis was computed using spiking responses during the delay and saccade epochs (one data point per trial). (D-F) Dynamics of slow drift for three example sessions when the SC slow drift axis was computed using spiking responses during the baseline, delay, visual and saccade epochs. Note that the session in (D) is the same as that show in (A-C). Baseline = 100ms before the onset of the target stimulus; Delay = 600 to 1100ms after the offset of the target stimulus; Stim. = 25ms to 125ms after the onset of the target stimulus; Sac. = 25ms before to 25ms after the onset of the saccade.

**Figure 2 - figure supplement 2.**
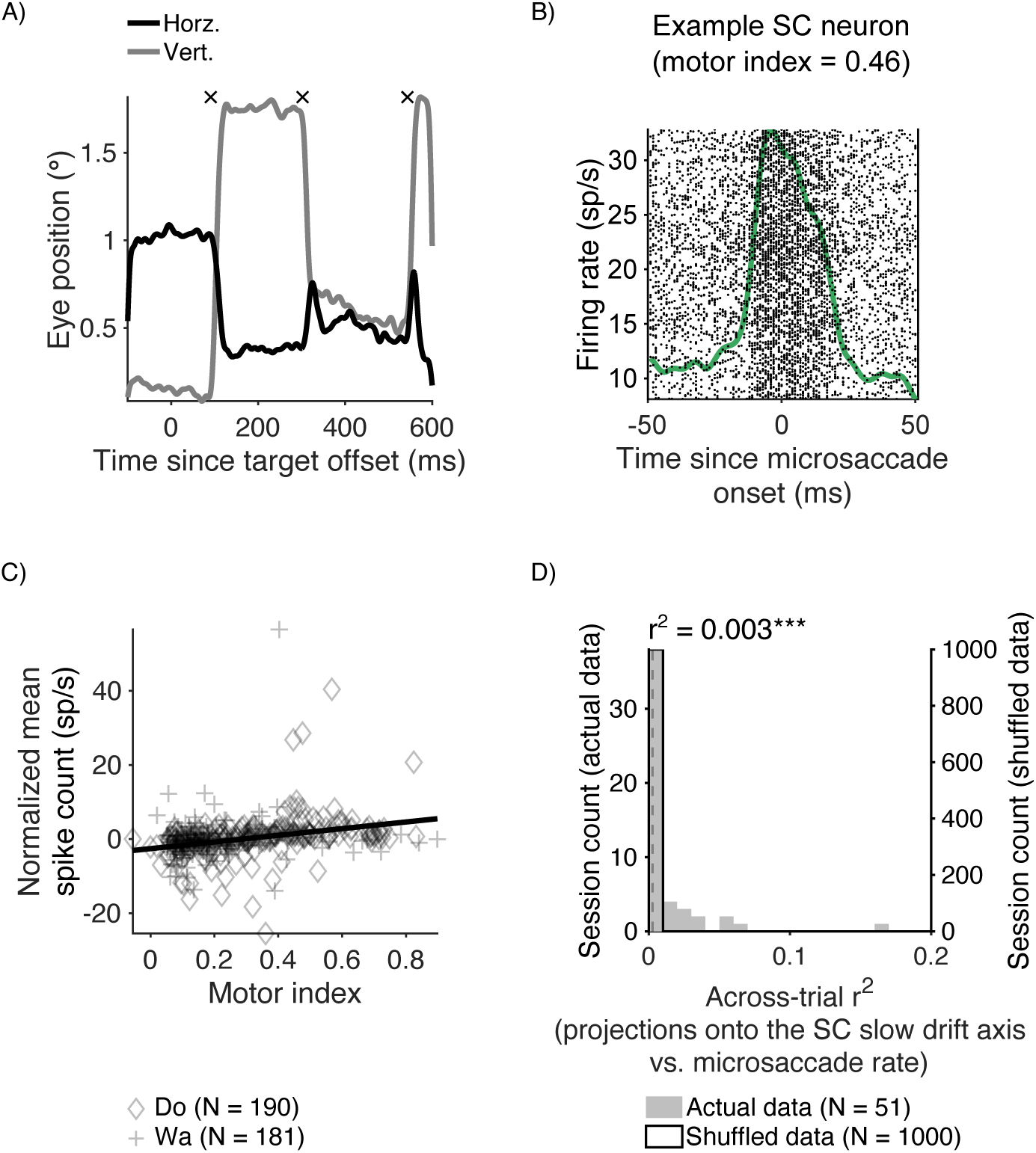
(A) Horizontal and vertical eye position on a single trial during the MGS task. Microsaccades were detected using the delay period using a velocity-base algorithm (Engbert and Kliegl, 2003) and are indicated by black crosses. (B) PSTH for an SC neuron that responded around the time of a microsaccade. Firing rates were computed in 1ms bins, averaged across trials and smoothed using a Gaussian function (σ = 5ms). Note that the targets were set to 3° in this session based on saccades evoked by microstimulation (see Methods). Previous research has shown that some SC neurons respond during microsaccades as well as to slightly larger saccades (Hafed and Krauzlis, 2012). This likely explains why this SC neuron, which had a RF at ∼3° based on saccades evoked by microstimulation, also responded around the time of a microsaccade. (C) Scatter plot showing the relationship between normalized mean firing rate in and around the time of a microsaccade and motor index for each recorded SC neuron. For each neuron, mean firing rates were computed in the period spanning 25ms before and 25m after a detected microsaccade, and were normalized by subtracting the mean activity across all trials during the same time period. This was done to control for the fact that some SC neurons (e.g. buildup neurons) have been found to increase their activity during the delay period on MGS tasks (Munoz and Wurtz, 1995). A multiple regression analysis was then performed to determine if normalized mean firing rate was associated with motor index. Results showed that the proportion of variance explained by the model, after controlling for possible intersubject differences, was 0.0791 (F(2, 368) = 15.8089; p < 0.001). Normalized mean firing rate was significantly associated with motor index such that it was higher for neurons that reside closer to the motor output (t = 5.3747; p < 0.001). (D) Histogram showing the proportion of variance explained between projections onto the SC slow drift axis and microsaccade rate. The median value across sessions is denoted by the dashed line. A p-value was computed by comparing the actual distribution of values to a null distribution (see Methods). p < 0.05*, p < 0.01**, p < 0.001***.

**Figure 5 - figure supplement 1.**
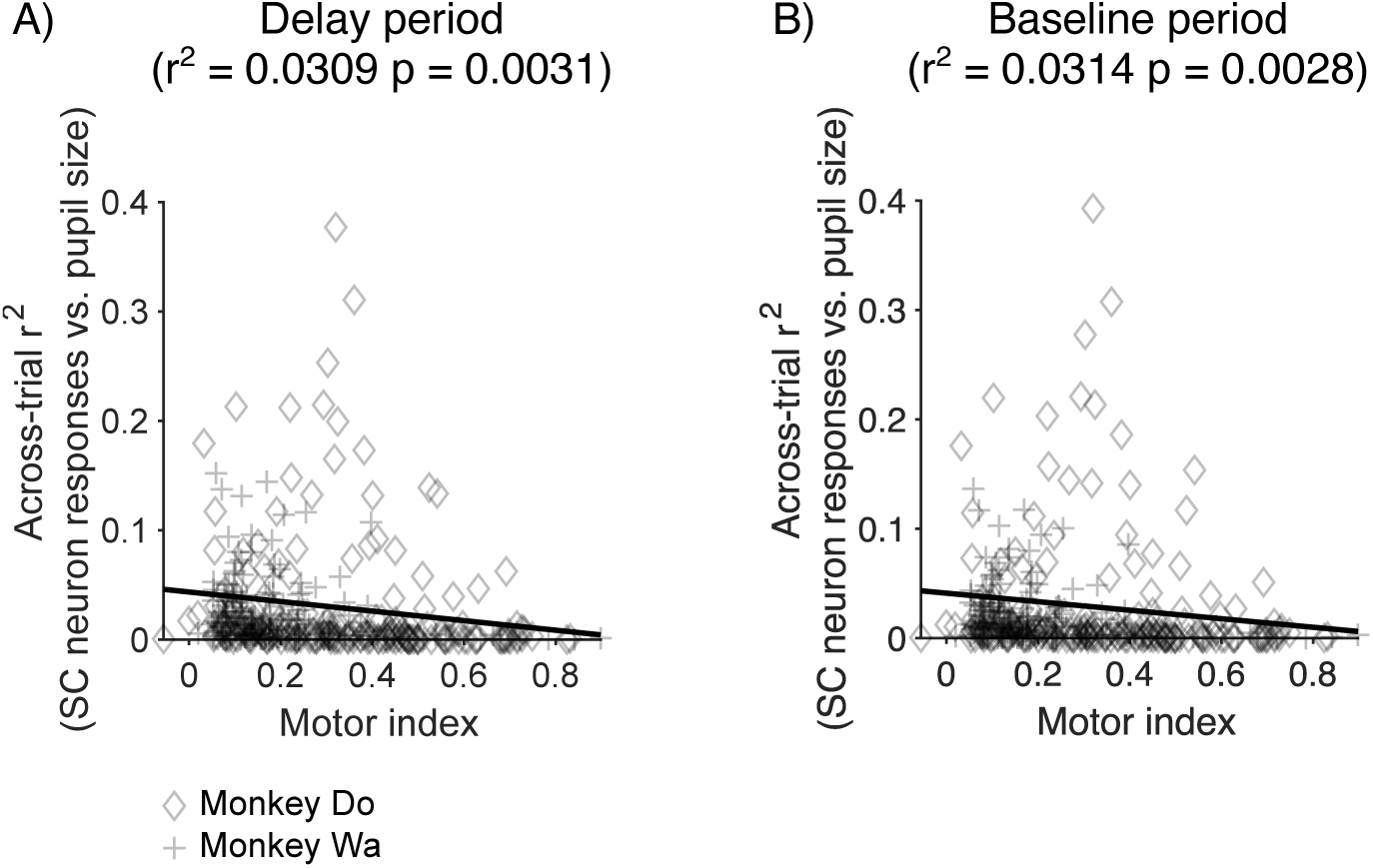
Scatter plot showing the strength of the relationship between individual SC neuron responses and pupil size. In (A) pupil size was computed during the delay period whereas in (B) it was computed during the baseline period.

## Notes

### Competing Interest Statement

The authors have declared no competing interest.

### Summary of Updates

This version contains revisions to the figures and text as a result of the review process.

## References

Allen KM, Lawlor J, Salles A, Moss CF (2021) Orienting our view of the superior colliculus: specializations and general functions. Curr Opin Neurobiol 71:119–126.

Ames KC, Churchland MM (2019) Motor cortex signals for each arm are mixed across hemispheres and neurons yet partitioned within the population response. Elife 8 Available at: 10.7554/eLife.46159.

Aoi MC, Mante V, Pillow JW (2020) Prefrontal cortex exhibits multidimensional dynamic encoding during decision-making. Nat Neurosci 23:1410–1420.

Aston-Jones G, Cohen JD (2005) An integrative theory of locus coeruleus-norepinephrine function: adaptive gain and optimal performance. Annu Rev Neurosci 28:403–450.

Ayar EC, Heusser MR, Bourrelly C, Gandhi NJ (2023) Distinct context- and content-dependent population codes in superior colliculus during sensation and action. Proc Natl Acad Sci U S A 120:e2303523120.

Basso MA, Bickford ME, Cang J (2021) Unraveling circuits of visual perception and cognition through the superior colliculus. Neuron 109:918–937.

Basso MA, May PJ (2017) Circuits for Action and Cognition: A View from the Superior Colliculus. Annu Rev Vis Sci 3:197–226.

Baumann MP, Bogadhi AR, Denninger AF, Hafed ZM (2023) Sensory tuning in neuronal movement commands. Proc Natl Acad Sci U S A 120:e2305759120.

Benavidez NL et al. (2021) Organization of the inputs and outputs of the mouse superior colliculus. Nat Commun 12:4004.

Bohlen MO, Warren S, May PJ (2017) A central mesencephalic reticular formation projection to medial rectus motoneurons supplying singly and multiply innervated extraocular muscle fibers. J Comp Neurol 525:2000–2018.

Bowers NR, Gautier J, Lin S, Roorda A (2021) Fixational eye movements in passive versus active sustained fixation tasks. J Vis 21:16.

Brainard D (1997) The Psychophysics Toolbox. Spat Vis 10:433–436.

Breton-Provencher V, Sur M (2019) Active control of arousal by a locus coeruleus GABAergic circuit. Nat Neurosci 22:218–228.

Cang J, Savier E, Barchini J, Liu X (2018) Visual Function, Organization, and Development of the Mouse Superior Colliculus. Annu Rev Vis Sci 4:239–262.

Castellote JM, Kumru H, Queralt A, Valls-Solé J (2007) A startle speeds up the execution of externally guided saccades. Exp Brain Res 177:129–136.

Cerkevich CM, Lyon DC, Balaram P, Kaas JH (2014) Distribution of cortical neurons projecting to the superior colliculus in macaque monkeys. Eye Brain 2014:121–137.

Chen C-Y, Hafed ZM (2018) Orientation and contrast tuning properties and temporal flicker fusion characteristics of primate superior colliculus neurons. Front Neural Circuits 12:58.

Churchland MM, Shenoy KV (2024) Preparatory activity and the expansive null-space. Nat Rev Neurosci 25:213–236.

Clark K, Squire RF, Merrikhi Y, Noudoost B (2015) Visual attention: Linking prefrontal sources to neuronal and behavioral correlates. Prog Neurobiol 132:59–80.

Cohen MR, Kohn A (2011) Measuring and interpreting neuronal correlations. Nat Neurosci 14:811–819.

Collins L, Francis J, Emanuel B, McCormick DA (2023) Cholinergic and noradrenergic axonal activity contains a behavioral-state signal that is coordinated across the dorsal cortex. Elife 12 Available at: 10.7554/eLife.81826.

Contadini-Wright C, Magami K, Mehta N, Chait M (2023) Pupil dilation and microsaccades provide complementary insights into the dynamics of arousal and instantaneous attention during effortful listening. J Neurosci 43:4856–4866.

Cowley BR, Snyder AC, Acar K, Williamson RC, Yu BM, Smith MA (2020) Slow Drift of Neural Activity as a Signature of Impulsivity in Macaque Visual and Prefrontal Cortex. Neuron 108:551–567.e8.

Crapse TB, Lau H, Basso MA (2018) A Role for the Superior Colliculus in Decision Criteria. Neuron 97:181–194.e6.

de Gee JW, Colizoli O, Kloosterman NA, Knapen T, Nieuwenhuis S, Donner TH (2017) Dynamic modulation of decision biases by brainstem arousal systems. Elife 6 Available at: 10.7554/eLife.23232.

Di Stasi LL, Catena A, Cañas JJ, Macknik SL, Martinez-Conde S (2013) Saccadic velocity as an arousal index in naturalistic tasks. Neurosci Biobehav Rev 37:968–975.

Ebitz RB, Hayden BY (2021) The population doctrine in cognitive neuroscience. Neuron 109:3055–3068.

Ebitz RB, Tu JC, Hayden BY (2020) Rules warp feature encoding in decision-making circuits. PLoS Biol 18:e3000951.

Edwards SB, Ginsburgh CL, Henkel CK, Stein BE (1979) Sources of subcortical projections to the superior colliculus in the cat. J Comp Neurol 184:309–329.

Elsayed GF, Lara AH, Kaufman MT, Churchland MM, Cunningham JP (2016) Reorganization between preparatory and movement population responses in motor cortex. Nat Commun 7:13239.

Engbert R, Kliegl R (2003) Microsaccades uncover the orientation of covert attention. Vision Res 43:1035–1045.

Gandhi NJ, Katnani HA (2011) Motor functions of the superior colliculus. Annu Rev Neurosci 34:205–231.

Goldberg ME, Wurtz RH (1972) Activity of superior colliculus in behaving monkey. I. Visual receptive fields of single neurons. J Neurophysiol 35:542–559.

Hafed ZM, Hoffmann K-P, Chen C-Y, Bogadhi AR (2023) Visual Functions of the Primate Superior Colliculus. Annu Rev Vis Sci 9:361–383.

Hafed ZM, Krauzlis RJ (2012) Similarity of superior colliculus involvement in microsaccade and saccade generation. J Neurophysiol 107:1904–1916.

Hafed ZM, Yoshida M, Tian X, Buonocore A, Malevich T (2021) Dissociable Cortical and Subcortical Mechanisms for Mediating the Influences of Visual Cues on Microsaccadic Eye Movements. Front Neural Circuits 15:638429.

Hennig JA, Oby ER, Golub MD, Bahureksa LA, Sadtler PT, Quick KM, Ryu SI, Tyler-Kabara EC, Batista AP, Chase SM, Yu BM (2021) Learning is shaped by abrupt changes in neural engagement. Nat Neurosci 24:727–736.

Heusser MR, Bourrelly C, Gandhi NJ (2022) Decoding the time course of spatial information from spiking and local field potential activities in the superior colliculus. eNeuro 9:ENEURO.0347-22.2022.

Hikosaka O, Wurtz RH (1983) Visual and oculomotor functions of monkey substantia nigra pars reticulata. III. Memory-contingent visual and saccade responses. J Neurophysiol 49:1268–1284.

Horwitz GD, Batista AP, Newsome WT (2004) Representation of an abstract perceptual decision in macaque superior colliculus. J Neurophysiol 91:2281–2296.

Huerta MF, Kaas JH (1990) Supplementary eye field as defined by intracortical microstimulation: connections in macaques. J Comp Neurol 293:299–330.

Ignashchenkova A, Dicke PW, Haarmeier T, Thier P (2004) Neuron-specific contribution of the superior colliculus to overt and covert shifts of attention. Nat Neurosci 7:56–64.

Isa T, Marquez-Legorreta E, Grillner S, Scott EK (2021) The tectum/superior colliculus as the vertebrate solution for spatial sensory integration and action. Curr Biol 31:R741–R762.

Issar D, Williamson RC, Khanna SB, Smith MA (2020) A neural network for online spike classification that improves decoding accuracy. J Neurophysiol 123:1472–1485.

Jagadisan UK, Gandhi NJ (2022) Population temporal structure supplements the rate code during sensorimotor transformations. Curr Biol 32:1010–1025.e9.

Johnston R, Snyder AC, Khanna SB, Issar D, Smith MA (2022a) The eyes reflect an internal cognitive state hidden in the population activity of cortical neurons. Cereb Cortex 32:3331–3346.

Johnston R, Snyder AC, Schibler RS, Smith MA (2022b) EEG Signals Index a Global Signature of Arousal Embedded in Neuronal Population Recordings. eNeuro 9 Available at: 10.1523/ENEURO.0012-22.2022.

Jolliffe IT, Cadima J (2016) Principal component analysis: a review and recent developments. Philos Trans A Math Phys Eng Sci 374:20150202.

Joshi S, Gold JI (2020) Pupil Size as a Window on Neural Substrates of Cognition. Trends Cogn Sci 24:466–480.

Joshi S, Gold JI (2022) Context-dependent relationships between locus coeruleus firing patterns and coordinated neural activity in the anterior cingulate cortex. Elife 11 Available at: 10.7554/eLife.63490.

Joshi S, Li Y, Kalwani RM, Gold JI (2016) Relationships between Pupil Diameter and Neuronal Activity in the Locus Coeruleus, Colliculi, and Cingulate Cortex. Neuron 89:221–234.

Jun EJ, Bautista AR, Nunez MD, Allen DC, Tak JH, Alvarez E, Basso MA (2021) Causal role for the primate superior colliculus in the computation of evidence for perceptual decisions. Nat Neurosci 24:1121–1131.

Kardamakis AA, Saitoh K, Grillner S (2015) Tectal microcircuit generating visual selection commands on gaze-controlling neurons. Proc Natl Acad Sci U S A 112:E1956–65.

Katnani HA, Gandhi NJ (2011) Order of operations for decoding superior colliculus activity for saccade generation. J Neurophysiol 106:1250–1259.

Katyal S, Zughni S, Greene C, Ress D (2010) Topography of covert visual attention in human superior colliculus. J Neurophysiol 104:3074–3083.

Kaufman MT, Churchland MM, Ryu SI, Shenoy KV (2014) Cortical activity in the null space: permitting preparation without movement. Nat Neurosci 17:440–448.

Kleiner M, Brainard D, Pelli D (2007) What’s new in psychtoolbox-3? perception 36. In: Ref Type: Abstract.

Knudsen EI (2011) Control from below: the role of a midbrain network in spatial attention. Eur J Neurosci 33:1961–1972.

Ko H-K, Poletti M, Rucci M (2010) Microsaccades precisely relocate gaze in a high visual acuity task. Nat Neurosci 13:1549–1553.

Komatsu H, Suzuki H (1985) Projections from the functional subdivisions of the frontal eye field to the superior colliculus in the monkey. Brain Res 327:324–327.

Krauzlis RJ, Lovejoy LP, Zénon A (2013) Superior colliculus and visual spatial attention. Annu Rev Neurosci 36:165–182.

Li L, Feng X, Zhou Z, Zhang H, Shi Q, Lei Z, Shen P, Yang Q, Zhao B, Chen S, Li L, Zhang Y, Wen P, Lu Z, Li X, Xu F, Wang L (2018) Stress Accelerates Defensive Responses to Looming in Mice and Involves a Locus Coeruleus-Superior Colliculus Projection. Curr Biol 28:859–871.e5.

Liu X, de Zwart JA, Schölvinck ML, Chang C, Ye FQ, Leopold DA, Duyn JH (2018) Subcortical evidence for a contribution of arousal to fMRI studies of brain activity. Nat Commun 9:395.

Mante V, Sussillo D, Shenoy KV, Newsome WT (2013) Context-dependent computation by recurrent dynamics in prefrontal cortex. Nature 503:78–84.

Massot C, Jagadisan UK, Gandhi NJ (2019) Sensorimotor transformation elicits systematic patterns of activity along the dorsoventral extent of the superior colliculus in the macaque monkey. Commun Biol 2:287.

Mathôt S (2018) Pupillometry: Psychology, Physiology, and Function. J Cogn 1:16.

May PJ (2006) The mammalian superior colliculus: laminar structure and connections. Prog Brain Res 151:321–378.

McCamy MB, Otero-Millan J, Leigh RJ, King SA, Schneider RM, Macknik SL, Martinez-Conde S (2015) Simultaneous recordings of human microsaccades and drifts with a contemporary video eye tracker and the search coil technique. PLoS One 10:e0128428.

Mohler CW, Wurtz RH (1976) Organization of monkey superior colliculus: intermediate layer cells discharging before eye movements. J Neurophysiol 39:722–744.

Müller JR, Philiastides MG, Newsome WT (2005) Microstimulation of the superior colliculus focuses attention without moving the eyes. Proceedings of the National Academy of Sciences 102:524–529.

Munoz DP, Wurtz RH (1995) Saccade-related activity in monkey superior colliculus. I. Characteristics of burst and buildup cells. J Neurophysiol 73:2313–2333.

Murphy PR, O’Connell RG, O’Sullivan M, Robertson IH, Balsters JH (2014) Pupil diameter covaries with BOLD activity in human locus coeruleus. Hum Brain Mapp 35:4140–4154.

Parthasarathy HB, Schall JD, Graybiel AM (1992) Distributed but convergent ordering of corticostriatal projections: analysis of the frontal eye field and the supplementary eye field in the macaque monkey. J Neurosci 12:4468–4488.

Pelli DG (1997) The VideoToolbox software for visual psychophysics: transforming numbers into movies. Spat Vis 10:437–442.

Perry VH, Cowey A (1984) Retinal ganglion cells that project to the superior colliculus and pretectum in the macaque monkey. Neuroscience 12:1125–1137.

Plummer NW, Chandler DJ, Powell JM, Scappini EL, Waterhouse BD, Jensen P (2020) An Intersectional Viral-Genetic Method for Fluorescent Tracing of Axon Collaterals Reveals Details of Noradrenergic Locus Coeruleus Structure. eNeuro 7 Available at: 10.1523/ENEURO.0010-20.2020.

Qu J, Zhou X, Zhu H, Cheng G, Ashwell KWS, Lu F (2006) Development of the human superior colliculus and the retinocollicular projection. Exp Eye Res 82:300–310.

Ratcliff R, Cherian A, Segraves M (2003) A comparison of macaque behavior and superior colliculus neuronal activity to predictions from models of two-choice decisions. J Neurophysiol 90:1392–1407.

Reimer J, McGinley MJ, Liu Y, Rodenkirch C, Wang Q, McCormick DA, Tolias AS (2016) Pupil fluctuations track rapid changes in adrenergic and cholinergic activity in cortex. Nat Commun 7:13289.

Robinson DA (1972) Eye movements evoked by collicular stimulation in the alert monkey. Vision Res 12:1795–1808.

Rosen MC, Freedman DJ (2025) Multiplexing of cognitive encoding by oculomotor networks leads to incidental gaze shifts. Proc Natl Acad Sci U S A 122:e2422331122.

Ruff DA, Cohen MR (2019) Simultaneous multi-area recordings suggest that attention improves performance by reshaping stimulus representations. Nat Neurosci 22:1669–1676.

Rust NC, Cohen MR (2022) Priority coding in the visual system. Nat Rev Neurosci 23:376–388.

Sara SJ, Bouret S (2012) Orienting and reorienting: the locus coeruleus mediates cognition through arousal. Neuron 76:130–141.

Semedo JD, Jasper AI, Zandvakili A, Krishna A, Aschner A, Machens CK, Kohn A, Yu BM (2022) Feedforward and feedback interactions between visual cortical areas use different population activity patterns. Nat Commun 13:1099.

Semedo JD, Zandvakili A, Machens CK, Yu BM, Kohn A (2019) Cortical Areas Interact through a Communication Subspace. Neuron 102:249–259.e4.

Setzer B, Fultz NE, Gomez DEP, Williams SD, Bonmassar G, Polimeni JR, Lewis LD (2022) A temporal sequence of thalamic activity unfolds at transitions in behavioral arousal state. Nat Commun 13:5442.

Smith MA, Sommer MA (2013) Spatial and temporal scales of neuronal correlation in visual area V4. J Neurosci 33:5422–5432.

Sparks DL (1986) Translation of sensory signals into commands for control of saccadic eye movements: role of primate superior colliculus. Physiol Rev 66:118–171.

Sparks DL, Hartwich-Young R (1989) The deep layers of the superior colliculus. Rev Oculomot Res 3:213–255.

Sparks DL, Mays LE (1990) Signal transformations required for the generation of saccadic eye movements. Annu Rev Neurosci 13:309–336.

Srinath R, Ruff DA, Cohen MR (2021) Attention improves information flow between neuronal populations without changing the communication subspace. Curr Biol 31:5299–5313.e4.

Stanton GB, Goldberg ME, Bruce CJ (1988) Frontal eye field efferents in the macaque monkey: II. Topography of terminal fields in midbrain and pons. J Comp Neurol 271:493–506.

Suzuki DG, Pérez-Fernández J, Wibble T, Kardamakis AA, Grillner S (2019) The role of the optic tectum for visually evoked orienting and evasive movements. Proc Natl Acad Sci U S A 116:15272–15281.

Turchi J, Chang C, Ye FQ, Russ BE, Yu DK, Cortes CR, Monosov IE, Duyn JH, Leopold DA (2018) The Basal Forebrain Regulates Global Resting-State fMRI Fluctuations. Neuron 97:940–952.e4.

van den Brink RL, Pfeffer T, Donner TH (2019) Brainstem Modulation of Large-Scale Intrinsic Cortical Activity Correlations. Front Hum Neurosci 13:340.

Wang C-A, Boehnke SE, Itti L, Munoz DP (2014) Transient pupil response is modulated by contrast-based saliency. J Neurosci 34:408–417.

Wang C-A, Boehnke SE, White BJ, Munoz DP (2012) Microstimulation of the monkey superior colliculus induces pupil dilation without evoking saccades. J Neurosci 32:3629–3636.

Wang C-A, Munoz DP (2018) Neural basis of location-specific pupil luminance modulation. Proc Natl Acad Sci U S A 115:10446–10451.

Wang C-A, Munoz DP (2021) Coordination of Pupil and Saccade Responses by the Superior Colliculus. J Cogn Neurosci 33:919–932.

Wheatcroft T, Saleem AB, Solomon SG (2022) Functional Organisation of the Mouse Superior Colliculus. Front Neural Circuits 16:792959.

White BJ, Berg DJ, Kan JY, Marino RA, Itti L, Munoz DP (2017) Superior colliculus neurons encode a visual saliency map during free viewing of natural dynamic video. Nat Commun 8:14263.

Winterson BJ, Collewijn H (1976) Microsaccades during finely guided visuomotor tasks. Vision Res 16:1387–1390.

Wurtz RH, Albano JE (1980) Visual-motor function of the primate superior colliculus. Annu Rev Neurosci 3:189–226.

Wylie DRW, Gutierrez-Ibanez C, Pakan JMP, Iwaniuk AN (2009) The optic tectum of birds: mapping our way to understanding visual processing. Can J Exp Psychol 63:328–338.

Zénon A, Krauzlis RJ (2012) Attention deficits without cortical neuronal deficits. Nature 489:434–437.

Zuber BL, Stark L, Cook G (1965) Microsaccades and the velocity-amplitude relationship for saccadic eye movements. Science 150:1459–1460.

